# Sas3-mediated histone acetylation regulates effector gene activation in a fungal plant pathogen

**DOI:** 10.1101/2023.01.18.524538

**Authors:** Marta Suarez-Fernandez, Rocío Álvarez-Aragón, Ana Pastor-Mediavilla, Alejandro Maestre-Guillén, Ivan del Olmo, Agustina De Francesco, Lukas Meile, Andrea Sánchez-Vallet

## Abstract

Effector proteins are secreted by plant pathogens to enable host colonization. Typically, effector genes are tightly regulated, have very low expression levels in axenic conditions, and are strongly induced during host colonization. Chromatin remodeling contributes to the activation of effector genes *in planta* by still poorly known mechanisms. In this work we investigated the role of histone acetylation in effector gene derepression in plant pathogens. We used *Zymoseptoria tritici*, a major pathogen of wheat, as a model to determine the role of lysine acetyltransferases (KATs) in plant infection. We showed that effector gene activation is associated with chromatin remodeling, featuring increased acetylation levels of histone H3 lysine 9 (H3K9) and 14 (H3K14) in effector loci. We functionally characterized the role of *Z. tritici* KATs and demonstrated their distinct contributions to growth, development, and infection. Sas3 is required for host colonization and pycnidia production, while Gcn5 has a major role in pycnidia production. Furthermore, we demonstrated that Sas3 is involved in acetylation of H3K9 and H3K14 in effector loci and in effector gene activation during plant infection. We propose that Sas3-mediated histone acetylation is required for spatiotemporal activation of effector genes and virulence of *Z. tritici*.

**IMPORTANCE:** Pathogen infections require the production of effectors that enable host colonization. Effectors have diverse functions and are only expressed at certain stages of the infection cycle. Thus, effector genes are tightly regulated by several mechanisms, including chromatin remodeling. Here, we investigate the role of histone acetylation in effector gene activation in the fungal wheat pathogen *Zymoseptoria tritici*. We demonstrated that lysine acetyltransferases (KATs) are essential for the spatiotemporal regulation of effector genes. We show that two KATs, Sas3 and Gcn5, are involved in leaf symptom development and pycnidia formation. Importantly, our results indicated that Sas3 controls histone acetylation of effector loci and is a regulator of effector gene activation during stomatal penetration. Overall, our work demonstrates the key role of histone acetylation in regulating gene expression associated with plant infection.

## 1. INTRODUCTION

Plant pathogens produce and secrete effectors into host tissues to facilitate colonization. Effectors have several functions including suppression of the immune response, alteration of plant development, acquisition of nutrients, and interference with the host microbiota (1). Effectors are frequently required at specific phases of the infection cycle (2). Consequently, the transcriptional control of effector genes is key to provide the pathogen with a dynamic infection machinery. Despite the importance of tight effector gene regulation in fungal plant pathogens, the mechanisms involved remain mostly enigmatic.

Chromatin remodeling is a pivotal mechanism of gene regulation and involves post-translational modifications of histone tails, such as acetylation and methylation. These modifications provide a conserved mechanism that modulates the accessibility of the transcription machinery to the DNA and thereby alters gene expression (3–5). Writing enzymes, including methylases, acetylases and erasing enzymes, such as demethylases and deacetylases, are dynamically involved in the posttranscriptional modification of histone tails in eukaryotes (6, 7). Effector genes are frequently located in heterochromatic regions of the genome (8, 9). In plant-associated fungi, including *Leptosphaeria maculans, Epichloë festucae, Magnaporthe oryzae* and *Zymoseptoria tritici*, effector genes are enriched in trimethylation of histone H3 lysine 9 (H3K9) and/or 27 (H3K27) in the absence of the host (10–13). During plant colonization, effector gene activation is associated with a tightly regulated reduction in the methylation levels in H3K9 and/or H3K27, as shown in *E. festucae* and *Z. tritici*. Accordingly, disruption of the key enzymes involved in methylation of H3K27 or H3K9 has been shown to enhance expression of effector genes and secondary metabolite gene clusters (10, 13). Thus, derepression of effector genes during host colonization involves changes in the chromatin state.

Although acetylation of specific residues of core histone tails has been shown to regulate transcription in eukaryotes (6, 7), the role of lysine acetyltransferases (KATs) in the overall fitness of fungal pathogens and in the spatiotemporal expression activation of effector genes remains to be largely understood. KATs transfer acetyl groups from acetyl-coenzyme A onto lysine residues of core histones and commonly form part of complexes (7). Frequently, KAT complexes harbor regulatory components that regulate KAT activity and substrate specificity to prevent uncontrolled histone acetylation (14). KATs are classified into different families, including the GNAT (from Gcn5-related N-acetyltransferase) and the MYST (MOZ, YBF2/SAS3, SAS2, and TIP60) families (7). Histone acetylation in filamentous fungi has been reported to regulate several biological processes such as growth, reproduction, secondary metabolite synthesis and pathogenicity. For instance, orthologues of Gcn5 mediate dimorphic changes, tolerance to stress, and virulence in *Ustilago maydis* (15), secondary metabolite regulation in *Aspergillus nidulans* (16), and stress tolerance and conidiation in *Alternaria alternata* (17). KATs from the MYST family are involved in growth and conidiation of *M. oryzae* and *A. alternata* (17, 18). In *Fusarium graminearum* KATs from the GNAT and MYST families mediate secondary metabolite regulation and virulence (19), highlighting the complexity of the role of KATs in trait regulation in fungi. Remarkably, histone acetylation is not only involved in fungus-plant interactions but also in fungus-bacterium interactions. Upon interaction of the filamentous fungus *A. nidulans* with the bacterium *Streptomyces rapamycinicus*, fungal secondary metabolite gene clusters are induced. This process involves acetylation of H3K9 and acetylation of histone H3 at lysine 14 (H3K14), and Gcn5 protein activity (16). Likewise, we hypothesized that histone acetylation plays a major role in effector gene activation in fungus-plant interactions. Given the important role of KATs in transcriptional activation in Eukaryotes and given the fact that effector genes are derepressed during host colonization (15–18), we propose that an increase of histone acetylation levels regulates activation of pathogen effector genes during plant infection.

*Z. tritici* is a major pathogen of wheat, causing significant yield losses in temperate climates (20). The infection cycle of *Z. tritici* initiates at the leaf surface with the germination of asexual or sexual spores. Emerged hyphae grow on the leaf surface and penetrate through the stomata. Subsequently, *Z. tritici* colonizes the apoplast and, after several days of infection, forms asexual fruiting bodies known as pycnidia (21). Chlorotic and necrotic symptoms are only observed after several days of infection of the pathogen, prior to asexual reproduction (22). *Z. tritici* mainly grows as a filamentous fungus on wheat leaf surfaces but can also grow as blastospores *in vitro* on rich media and occasionally on the leaf surface (23). Various effector genes are strongly induced during plant infection at different stages of the *Z. tritici* life cycle, *AvrStb6* and *Avr3D1* being activated during stomatal penetration and apoplast colonization, while *Mycgr3G76589* is expressed at later stages of the infection life cycle (10, 24, 25). Promoter activity and local reduction of histone methylation levels are required for the specific expression pattern of effector genes (10). Our integrative study aimed to determine the role of histone acetylation and KATs in effector gene regulation in *Z. tritici*. We demonstrated that dynamic histone acetylation of H3K9 and H3K14 is associated with expression activation of effector genes and host colonization.

## 2. RESULTS

### 2.1 *Z. tritici* has 7 orthologues of KATs

We first aimed to identify KAT orthologues in *Z. tritici*. To achieve this, we performed a BLAST search on annotated *Z. tritici* genes using previously characterized *Saccharomyces cerevisiae* KATs as queries. In addition, we used the dbHiMo database (26), which comprises histone-modifying enzymes from several fungal species including *Z. tritici*. A reverse BLAST analysis with the identified putative *Z. tritici* KAT orthologues was subsequently performed on the *S. cerevisiae* genome. We found 3 KAT orthologues from the MYST family, 2 from the GNAT family, and 1 from the specific fungal family RTT109 (Table 1). Additionally, 1 orthologue of Gcn5-related N-acetyltransferase (Ngs1; Table 1) previously identified in *Candida albicans* was also identified in *Z. tritici* (27).

**Table 1.** Classification of lysine acetyltransferase (KAT) orthologues in *Zymoseptoria tritici* strain ST99CH_3D7.

| KAT<br>nomenclature | Previous<br>nomenclature | KAT family | Gene ID in *Zymoseptoria*<br>*tritici* strain ST99CH_3D7 |
| --- | --- | --- | --- |
| KAT1 | Hat1 | GNAT | Not found |
| KAT2 | Gcn5 | GNAT | *3D7.g4775* |
| KAT5 | Esa1 | MYST | *3D7.g9281* |
| KAT6 | Sas3 | MYST | *3D7.g4263* |
| KAT8 | Sas2 | MYST | *3D7.g7031* |
| KAT9 | Elp3 | GNAT | *3D7.g8500* |
| KAT11 | Rtt109 | RTT109 | *3D7.g8867* |
| - | Ngs1 | GNAT | *3D7.g2851* |

The BLASTp (Table S1) and dbHiMo analyses provided a consistent classification of 3D7.g7031 as Sas2 (KAT8) and 3D7.g9281 as Esa1 (KAT5). However, the classification of 3D7.g4263 was conflicting since dbHiMo predicted it to be Esa1, while the BLASTp analysis classified it as a Sas3 protein. To properly classify KAT orthologues, a phylogenetic tree was constructed using KATs of the MYST family from different fungal species. The phylogenetic analysis showed that 3D7.g4263 clusters with Sas3 (KAT6) protein orthologues, that 3D7.g7031 is a Sas2 orthologue, and that 3D7.g9281 is an Esa1 orthologue (Figure 1A; Table S2; Table 1). We identified the expected MOZ/SAS domain and the MYST family zinc finger domain in the *Z. tritici* orthologues of Esa1 and Sas2 using HMMER (Figure 1C). In addition to these two domains, Esa1 harbors an RNA-binding domain near the N-terminus. Sas3 is the largest protein identified and contains two MOZ/SAS domains next to a MYST family zinc finger domain and a plant homeodomain (PHD) - finger domain.

**Figure 1.**
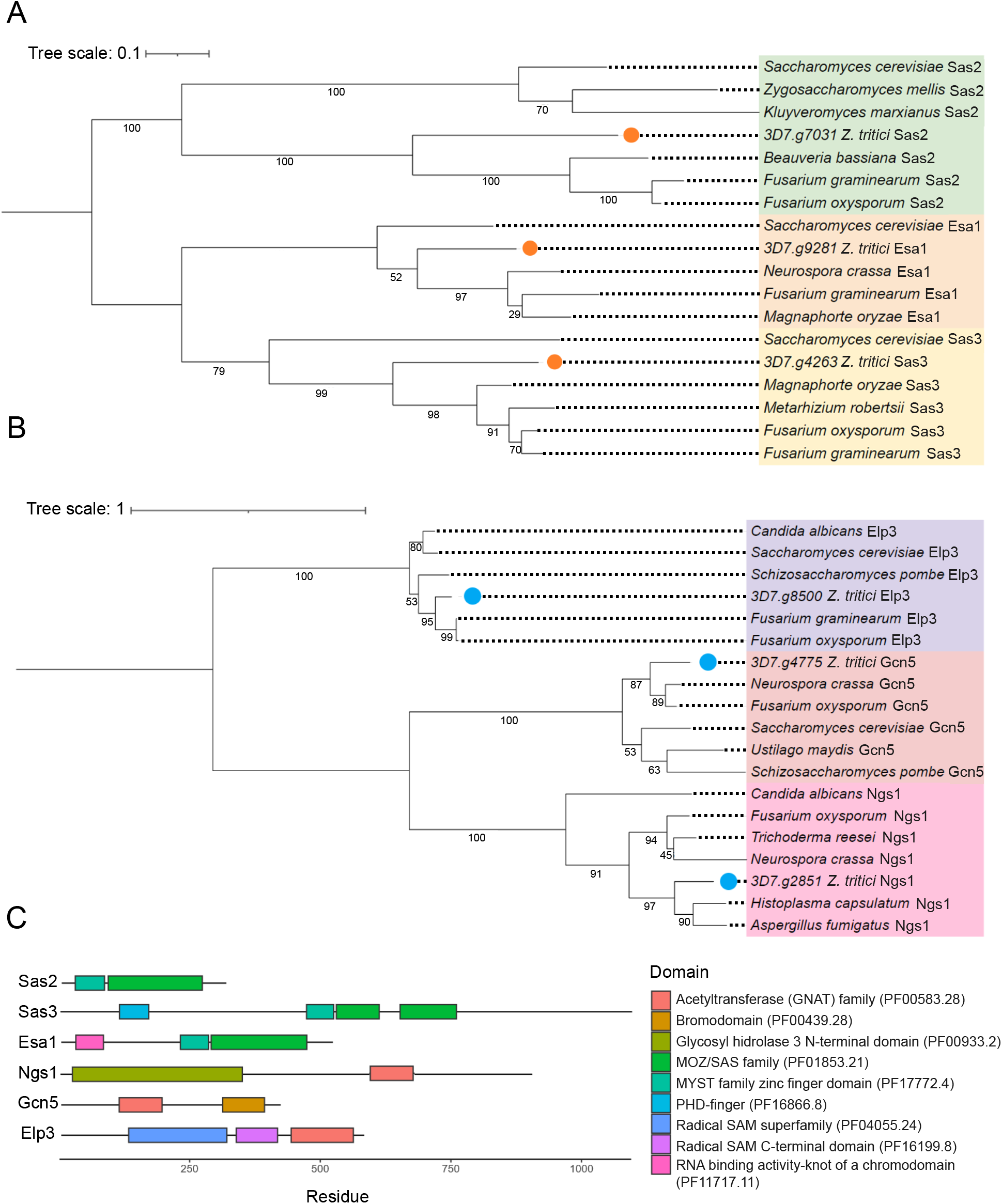
*Zymoseptoria tritici* has 3 lysine acetyltransferase (KAT) orthologues belonging to the MYST family (Esa1, Sas2 and Sas3) and 3 belonging to the GNAT family (Ngs1, Gcn5 and Elp3). A) Phylogenetic tree of the MYST family protein members from different fungal organisms. MYST sequences belonging to *Z. tritici* are indicated with orange dots. The protein names are colored according to their classification: Sas2 (KAT8; green), Esa1 (KAT5; orange), and Sas3 (KAT6; yellow). B) Phylogenetic tree of GNAT family proteins from different fungal organisms. GNAT sequences belonging to *Z. tritici* are indicated with blue dots. The protein names are colored according to the type of enzyme: Elp3 (KAT9; purple), Gcn5 (KAT2; light red), and Ngs1 (pink). The numbers below the branches represent the support values from 1000 bootstrap replicates using the maximum likelihood method. Trees have been rooted using the midpoint root method. Tree scale indicates branch length in the tree. Units are given in residue substitution per site. C) Domains identified in the KAT proteins of *Z. tritici*.

The phylogenetic analysis of KATs of the GNAT family indicated that 3D7.g2851, 3D7.g4775 and 3D7.g8500 cluster with Ngs1, Gcn5 (KAT2) and Elp3 (KAT9) proteins, respectively (Figure 1B; Table S2; Table 1). All the identified KAT orthologues belonging to the GNAT family contain a GNAT acetyltransferase domain. Gcn5 additionally has a bromodomain in its C-terminal region, while the Elp3 orthologue contains 2 radical SAM domains. Ngs1 contains a glycosyl hydrolase family 3 (GH3) domain (Figure 1C), as previously described for orthologues of this KAT in other organisms (27).

### 2.2 KAT genes in *Z. tritici* are differentially expressed during plant infection

We hypothesized that KATs involved in effector gene regulation might be expressed during plant infection and they might exhibit a similar expression pattern as effector genes. Therefore, we performed a transcriptomic analysis of the genes encoding the identified KATs and compared them with the expression of three effector genes formerly shown to be epigenetically regulated (10): *Avr3D1* (24), *AvrStb6* (25) and *Mycgr3G76589* (28). For this purpose, we used data from previously published RNA-seq studies (23, 29). The MYST family orthologues (*Esa1, Sas2* and *Sas3*) were expressed during host colonization, displaying low expression levels at the beginning of the infection and a peak of expression at 12-14 days post-infection (dpi). The three GNAT family members were also expressed during host colonization, *Elp3* exhibiting the lowest expression levels at 14 dpi (Figure 2). Based on the different gene expression patterns of the KAT members, we hypothesized that they might have distinct roles in growth, development, and virulence.

**Figure 2.**
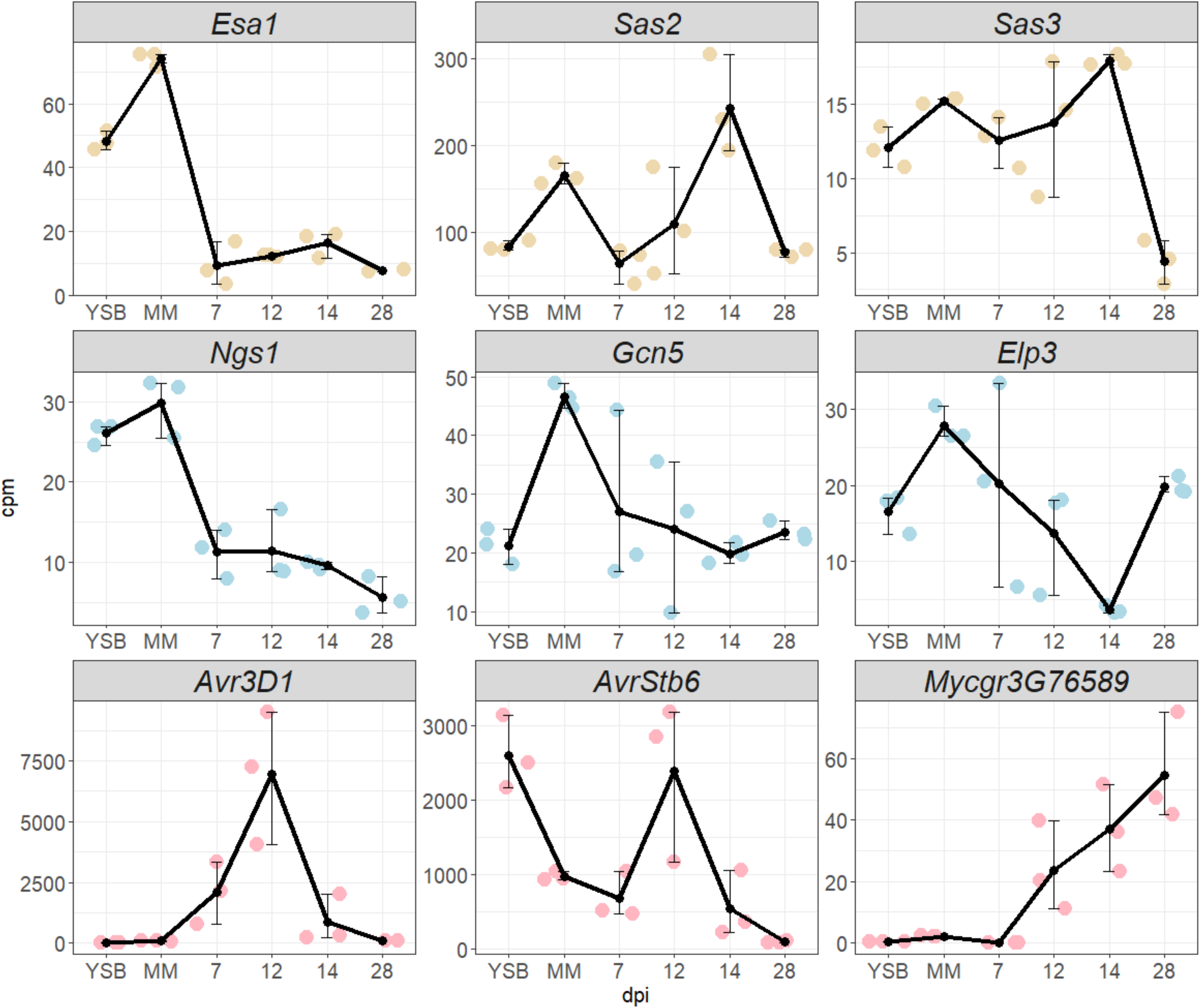
Specific regulation of *Z. tritici* lysine acetyltransferase (KAT) genes during infection. Expression levels of the KAT genes *Esa1* (MYST family), *Sas2* (MYST family), *Sas3* (MYST family), *Ngs1* (GNAT family), *Gcn5* (GNAT family), *Elp3* (GNAT family) and the effector genes *Avr3D1, AvrStb6* and *Mycgr3G76589* under axenic conditions in two media with different nutrient content: Yeast extract sucrose broth (YSB) and minimal medium (MM); and during infection at different time points (7-, 12-, 14- and 28-days post infection; dpi). Data were obtained from previously published RNA-seq studies (NCBI accessions: SRA SRP152081 and SRP077418) (23, 29). cpm: counts per million mapped reads.

### 2.3 Histone acetylation levels in effector loci increase during plant infection

To evaluate the changes in acetylation of H3K9 and H3K14 during plant infection in effector loci, we performed a chromatin immunoprecipitation assay followed by quantitative PCR (ChIP-qPCR). We infected wheat plants with *Z. tritici* and harvested the second leave at 11 dpi, which is approximately the time point at which maximum levels of effector transcripts can be observed (Figure 2; (10)). We additionally analyzed histone acetylation in *Z. tritici* grown under axenic conditions. We evaluated the acetylation levels of H3K9 and H3K14 in different regions of *AvrStb6*: 1000 base pairs (bp; −1000), 500 bp (−500), 300 bp (−300) and 50 bp (−50) upstream of the start codon, and within the open reading frame region (ORF). We also evaluated the acetylation of these two marks 300 bp upstream of the start codon of *Avr3D1*. A TFIIIC transcription factor complex unit (*3D7.g8520; TFIIIC*) was used as control. As expected (30, 31), the acetylation levels of the control and *AvrStb6* (−1000) remained stable *in planta* compared to axenic conditions. We observed an increase in the acetylation levels of H3K9 and H3K14 at the loci of the effector genes *Avr3D1* and *AvrStb6* (Figure 3). The ChIP-qPCR results support a possible role of histone acetylation in the activation of effector genes during infection.

**Figure 3.**
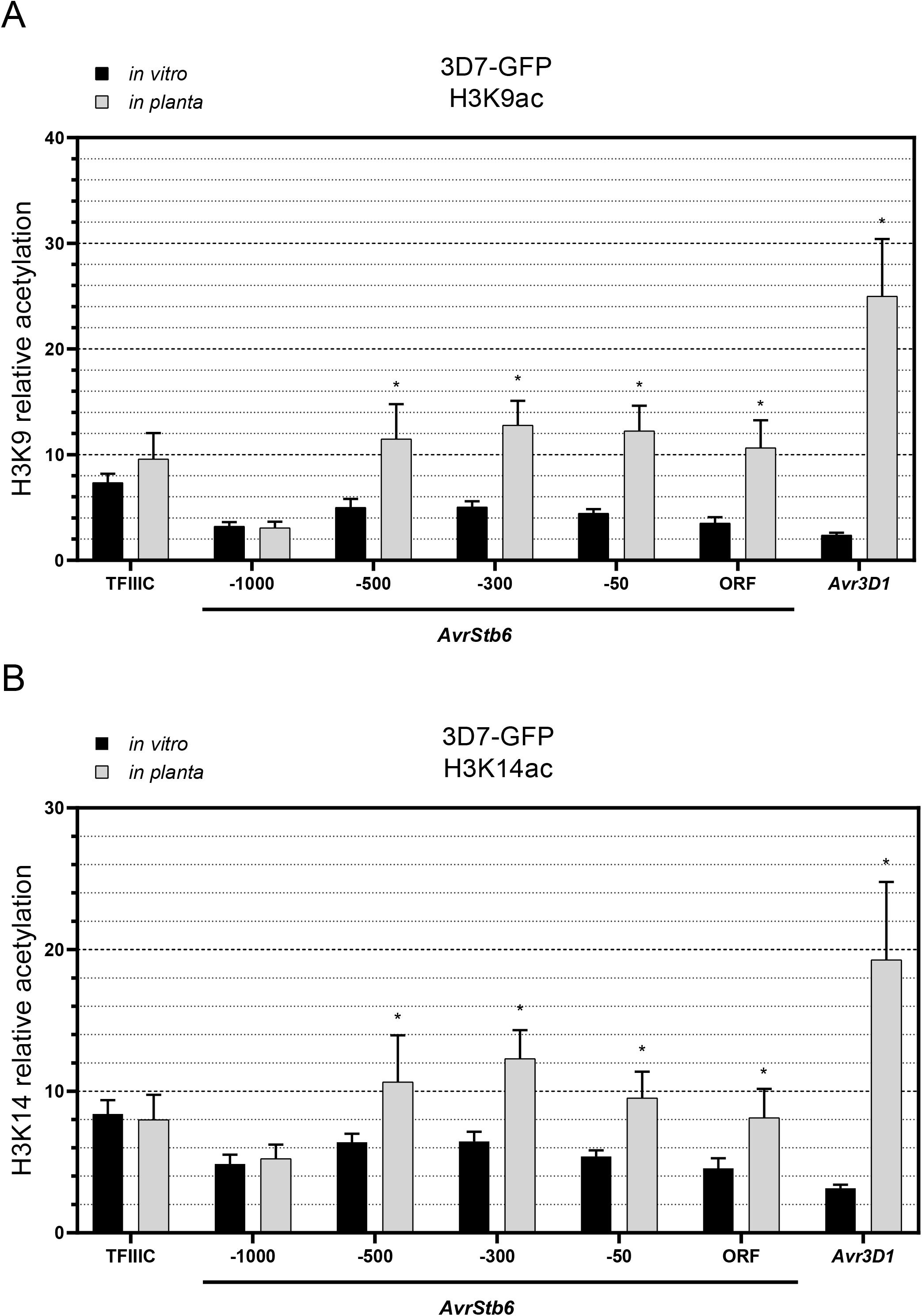
Acetylation levels of histone H3 lysine 9 (H3K9) and 14 (H3K14) in *Z. tritici* increase during plant infection. Relative acetylation of H3K9 (A) and H3K14 (B) in 3D7-GFP *in vitro* and *in planta* in different regions of *AvrStb6*: 1000 base pairs (bp; −1000), 500 base pairs (−500), 300 bp (−300) and 50 bp (−50) upstream of the start codon, within the open reading frame (ORF), and 300 bp upstream of the start codon of *Avr3D1. TFIIIC* (100 bp upstream of start codon) was used as controls. Chromatin immunoprecipitation experiments were performed during host colonization at 11 days post infection (dpi). Acetylation levels are shown relative to H3 levels. Bars show the average of three independent biological replicates and the error bars represent the standard error of the mean. Asterisks indicate significant differences between infection and axenic conditions according to two-way ANOVA and Bonferroni tests (p < 0.05).

### 2.4 KAT orthologues in *Z. tritici* are involved in growth and colony development

To determine the function of the KAT orthologues of *Z. tritici* in development and virulence, we obtained loss-of-function mutants in the genes *Sas2*, *Sas3*, *Ngs1, Gcn5* and *Elp3*. We first determined the role of the investigated KATs in development. We measured the area of colonies of the KAT mutants grown on yeast-malt-sucrose agar (YMA; Figure S1). Δ*Sas3* and Δ*Gcn5* colonies were significantly smaller than colonies of the control. Interestingly, Δ*Sas2* lines displayed the opposite phenotype, with larger colony diameters than the controls, most likely due to their hyphal-like growth, as observed on the colony edges (Figure S1). We therefore suggest that Sas2, Sas3 and Gcn5 might be involved in growth and/or development, with Sas2 probably being a negative regulator of growth and hyphal switch.

We addressed the role of the KATs in stress tolerance by assessing the performance of the mutants under different stresses, including high temperature (28°C), salt (NaCl; 0.5 M), H_2_O_2_ (1 mM), osmotic (sorbitol; 1 M) and cell wall (Calcofluor white; 200 ng·μL^−1^, and Congo red; 2 mg·mL^−1^) stresses on YMA. Additionally, we quantified growth in the presence of different carbon sources, such as fructose (5 g·L^−1^), galactose (50 mM), N-acetylglucosamine (GlcNAc, 2.5 mM), and glucose (2.5 mM) in a nutrient poor minimal medium (MM). Colony development of the mutants was compared with growth under standard conditions (YMA 18°C) and to the control line (3D7-GFP). Δ*Sas3* colonies were smaller than those of the control line under all the conditions, supporting the role of Sas3 in growth and/or development. The other tested mutants grew similarly to 3D7-GFP under all the stress conditions, except for Δ*Gcn5* and Δ*Ngs1*, which were slightly more resistant to Congo red than the control line (Figure S2). Therefore, we concluded that *Z. tritici* KATs are not positive regulators of stress tolerance.

### 2.5 Sas3 and Gcn5 are involved in virulence

We further investigated the role of KATs in host colonization on the susceptible wheat cultivar Runal (Figure 4; Figures S3 and S4). All the KAT mutants developed a similar biomass to that of the control at 10 dpi, except for Δ*Sas3* and Δ*Ngs1*, which had a significantly lower biomass (Figure 4A). However, at shorter time points (6 dpi), Δ*Sas3* grew to similar levels as the control (Figure 4B; Figure S5). In accordance with the reduction in biomass, Δ*Sas3* developed less disease symptoms, as determined by the percentage of leaf area covered by lesions (PLACL), and less pycnidia (Figure 4; Figures S3 and S6). Additionally, we observed a slightly faster production of symptoms by the mutant Δ*Gcn5* (Figure 4D; Figures S4 and S6). Nevertheless, this faster spread of disease symptoms did not lead to a higher production of pycnidia. Instead, we observed that Δ*Gcn5* generated very few pycnidia (Figure 4E; Figures S4, S6 and S7). Δ*Elp3* showed a slight reduction in pycnidia production per leaf lesion at 20 dpi (Figure 4E; Figure S4), suggesting a contribution of Elp3 to asexual reproduction. Additionally, Δ*Sas2* showed a slightly altered infection phenotype, manifested by red/orange spots on infected leaves, but no differences in PLACL or in pycnidia production were detected (Figure 4; Figures S3 and S6). Overall, *Gcn5* and *Sas3* disruption led to the highest effect on virulence and/or pycnidia production. To confirm that the observed phenotype of Δ*Gcn5* and Δ*Sas3* was due to the disruption of the KAT genes, we obtained complementation lines which expressed the wildtype version of *Gcn5* and *Sas3* in the mutant backgrounds. We observed a restoration of virulence and pycnidia production in both complementation lines (Figure S7). These results demonstrate that Sas3 is involved in virulence and that Sas3 and Gcn5 mediate pycnidia formation, indicating the key role of *Z. tritici* KATs in virulence and reproduction.

**Figure 4.**
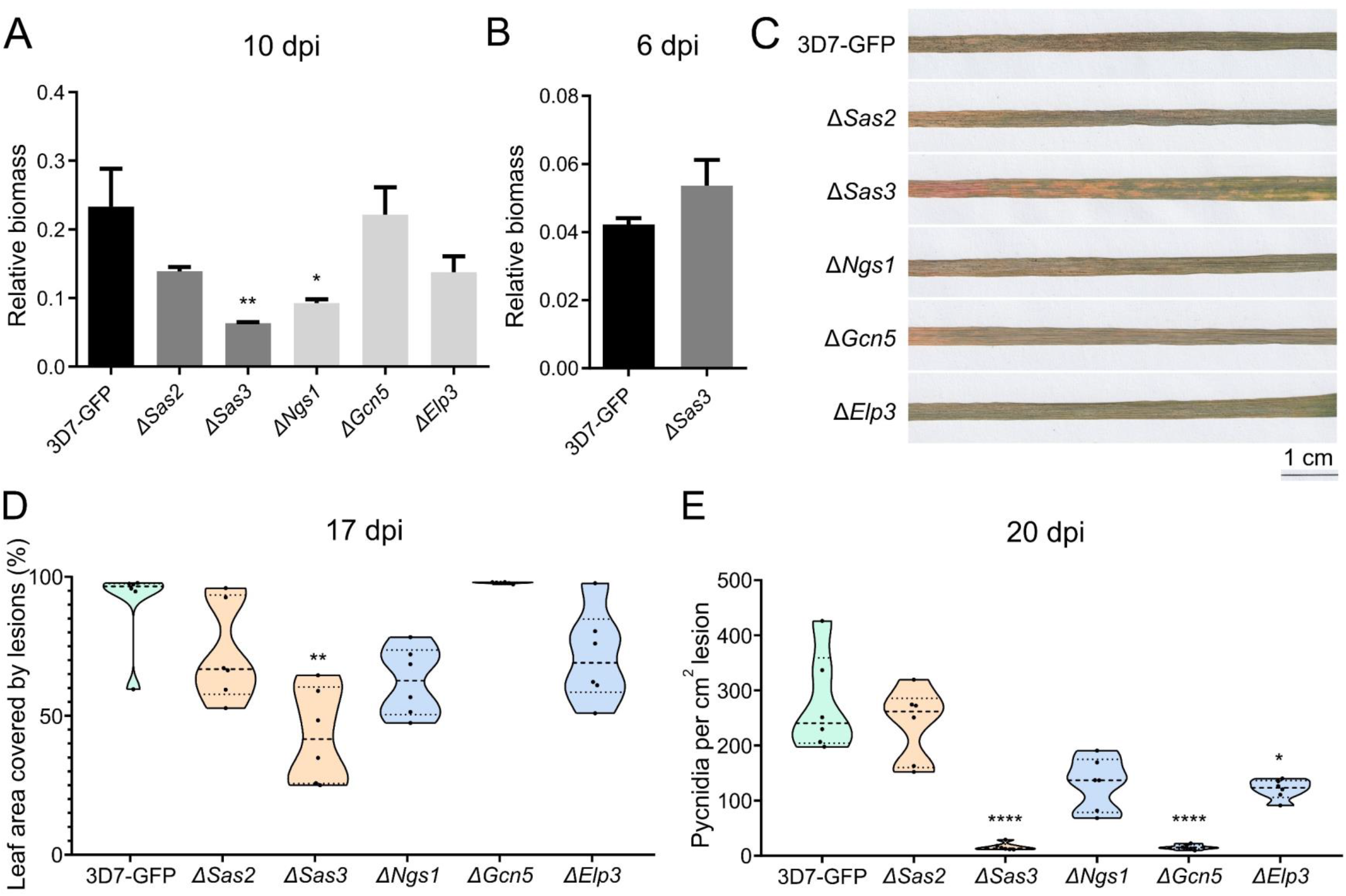
Lysine acetyltransferases (KATs) are involved in *Z. tritici* infection. Relative biomass to a reference control of the control (3D7-GFP) and the KAT mutants (Δ*Ngs1*, Δ*Sas2*, Δ*Sas3*, Δ*Gcn5* and Δ*Elp3*) during infection at 10 days post infection (dpi) (A) and of 3D7-GFP and Δ*Sas3* at 6 dpi (B). Representative pictures of wheat leaves infected with 3D7-GFP and the knockout mutants in the KAT genes at 20 dpi (C). Percentage of leaf area covered by lesions (PLACL) at 17 dpi (D) and pycnidia per cm^2^ of lesion at 20 dpi (E) of wheat plants infected with the control and the KAT mutants. In A and B, black bars represent 3D7-GFP, dark grey bars represent MYST-family mutants and light grey bars represent GNAT-family mutants. In D and E, green represents 3D7-GFP, orange represents MYST-family mutants and blue represents GNAT-family mutants. Dashed lines represent the median, dotted lines represent first and third quartiles and black dots represent individual data points. Asterisks indicate significant differences with 3D7-GFP according to Kruskal-Wallis and Dunn’s tests (* p < 0.05; ** p < 0.01; **** p < 0.0001).

### 2.6 KAT mutants are impaired in expression regulation of effector genes under axenic conditions

We next determined the role of *Z. tritici* KATs in regulation of the infection machinery by analyzing the expression levels of the effector genes *Avr3D1, AvrStb6* and *Mycgr3G76589* by qRT-PCR under axenic conditions. The expression of *Avr3D1* was drastically reduced in Δ*Sas3* (Figure 5A), and the expression of *AvrStb6* was reduced in Δ*Sas2*, Δ*Sas3* and Δ*Gcn5* (Figure 5B) under axenic conditions. *Mycgr3G76589* expression was unchanged in the mutants (Figure S8).

**Figure 5.**
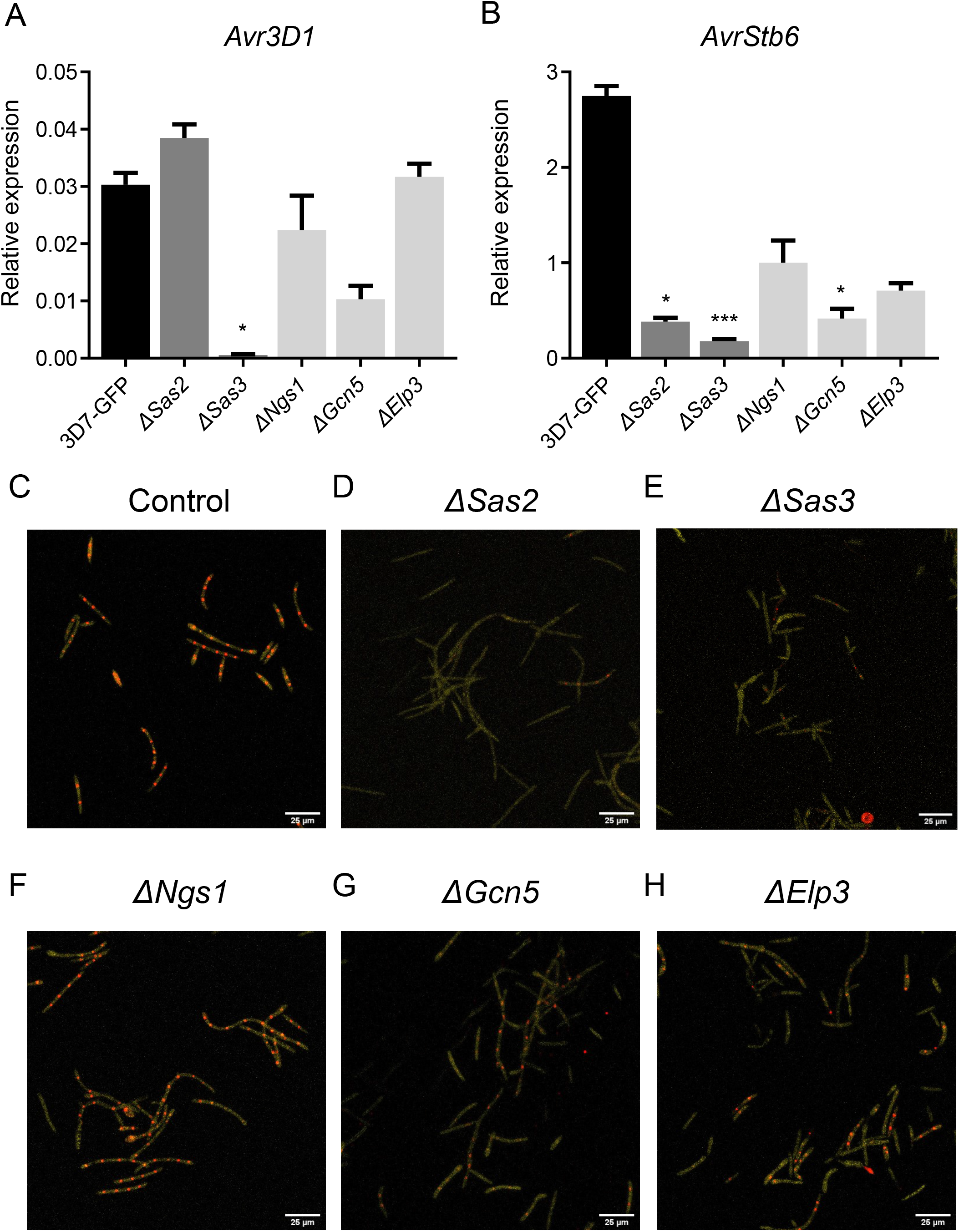
Lysine acetyltransferases (KATs) regulate effector gene expression under axenic conditions. Relative expression of *Avr3D1* (A) and *AvrStb6* (B) in the control (3D7-GFP) and the KAT mutants grown on yeast-malt-sucrose agar (YMA) for 6 days. *β-tubulin* and *histone H3* were used both as reference genes. Each bar corresponds to the mean expression value of 3 biological replicates and error bars represent the standard error of the mean. Asterisks indicate statistical differences with 3D7-GFP according to Kruskal-Wallis with uncorrected Dunn’s tests. (* p < 0.05; *** p < 0.001). Expression pattern of *AvrStb6* at the cellular level at 6 days post infection (dpi) in the control reporter line (C); and the control reporter line lacking *Sas2* (D); *Sas3* (E); *Ngs1* (F); *Gcn5* (G); and *Elp3* (H). In the reporter line, mCherry fused to histone 1 was expressed under the control of the *AvrStb6* promoter in the *AvrStb6* locus. This allowed the localization of the reporter to the nucleus (red dots) and therefore to monitor the activity of the *AvrStb6* promoter at the single-cell level (10). Fungal blastospores are labelled with mTurquoise2 and shown in yellow. Calibration bars correspond to 25 μm.

To determine the role of the investigated KATs in effector gene regulation at the cellular level, we disrupted the KAT genes in a reporter line that harbors His1-mCherry located at the *AvrStb6* locus, and under the control of the *AvrStb6* promoter (10). The fusion of histone 1 (His1) with mCherry enabled its nuclear localization and allowed monitoring *AvrStb6* expression at the cellular level. In the control reporter line growing under axenic conditions, mCherry was detected, indicating that the *AvrStb6* promoter is partially active when *Z. tritici* grows in the absence of the host (Figures 5C and 2). Δ*Ngs1* and Δ*Elp3* showed the same expression pattern as the control (Figures 5F and 5H), while Δ*Sas2*, Δ*Sas3* and Δ*Gcn5* showed a decrease in the levels of mCherry accumulation under axenic conditions (Figures 5D, 5E and 5G). These results confirmed that *Z. tritici* Sas2, Sas3 and Gcn5 are involved in effector gene regulation under axenic conditions.

### 2.7 Effector gene expression is altered during plant infection in the KAT mutants

Effector genes are key for plant colonization and highly induced during infection (Figure 2). We hypothesized that histone acetylation might be required for effector gene upregulation. We determined the expression levels of 4 effector genes (*Avr3D1 (3D7.g7883* reannotated in (24)), *AvrStb6 (3D7.g5586), AvrStb9 (3D7.g741*; (32)), and *Mycgr3G76589 (3D7.g10118)*) by qRT-PCR during plant infection in KAT knockouts (Figure 6). The expression levels of *Avr3D1* and *AvrStb6* were higher in Δ*Ngs1* and Δ*Gcn5*, while *AvrStb6* expression was reduced in Δ*Sas2* and Δ*Sas3* at 10 dpi (Figures 6A and 6B). We also observed reduced expression of *Mycgr3G76589* in all mutants except Δ*Gcn5* (Figure 6D) and a reduction of *AvrStb9* expression in Δ*Sas3* (Figure 6C). These results suggest that KATs are involved in infection and in the proper regulation of effector gene expression during early stages of plant colonization.

**Figure 6.**
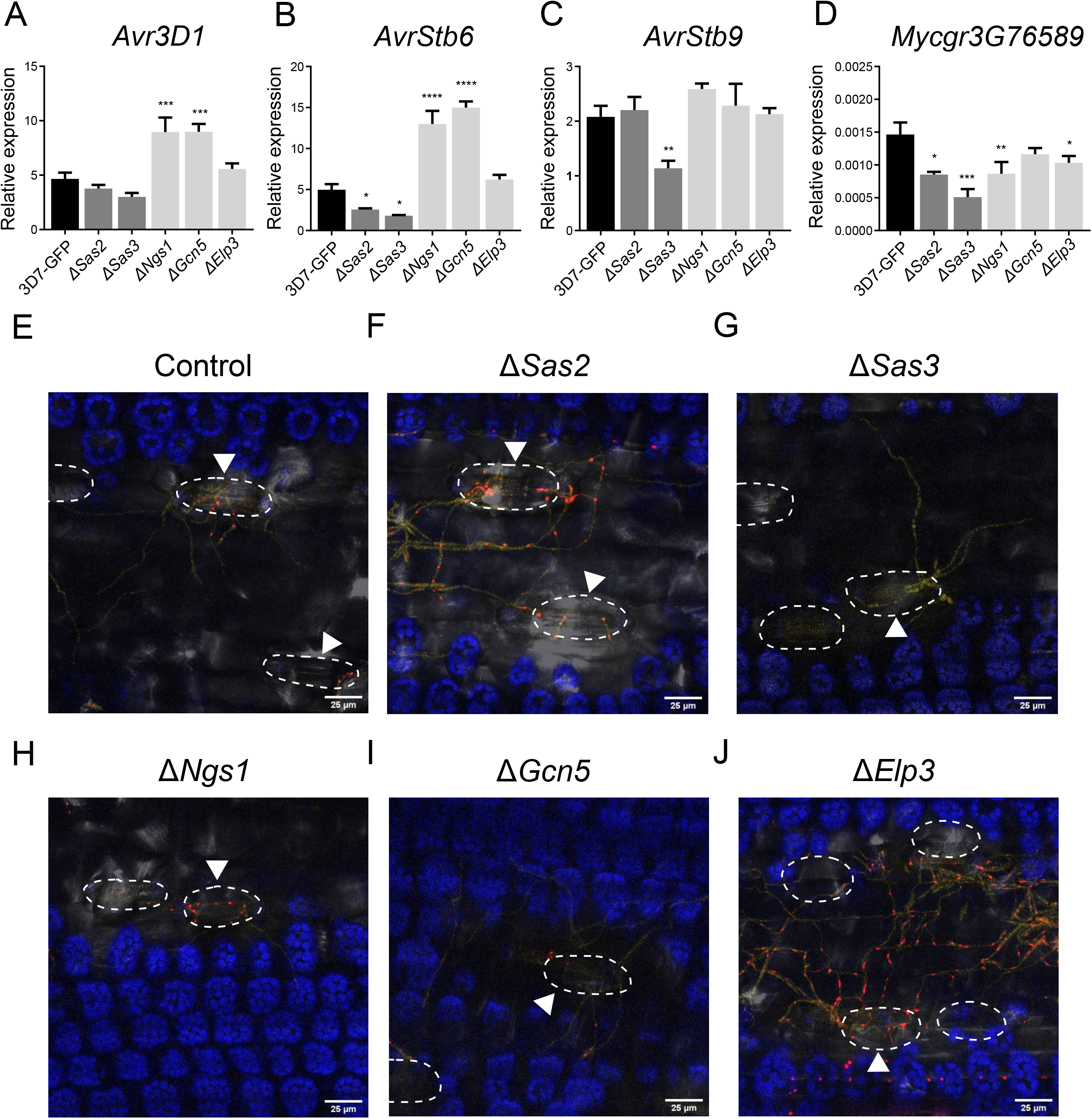
Lysine acetyltransferases (KATs) are involved in effector gene regulation during plant colonization. Relative expression of *Avr3D1* (A), *AvrStb6* (B), *AvrStb9* (C) and *Mycgr3G76589* (D) in the control (3D7-GFP) and KAT mutants (Δ*Ngs1*, Δ*Sas2*, Δ*Sas3*, Δ*Gcn5* and Δ*Elp3*) during wheat infection at 10 days post infection (dpi). *β-tubulin* and *histone H3* were used both as reference genes. Bars correspond to the mean expression value of 3 biological replicates per treatment and error bars represent standard error of the mean. Asterisks indicate significant differences with 3D7-GFP according to Kruskal-Wallis and Dunn’s tests (* p < 0.05; ** p < 0.01). Expression pattern of *AvrStb6* at the cellular level at 6 dpi in the E) control reporter line; and the control reporter line lacking *Sas2* (F); *Sas3* (G); *Ngs1* (H); *Gcn5* (I); and *Elp3* (J). In the reporter line, mCherry fused to histone 1 was expressed under the control of the *AvrStb6* promoter in the *AvrStb6* locus. This allowed the localization of the reporter to the nucleus (red dots) and therefore monitoring the activity of the *AvrStb6* promoter at the single-cell level (10). Fungal hyphae are labelled with mTurquoise2 and shown in yellow. Chloroplasts are indicated in blue. White discontinuous lines indicate the stomata. Hyphae penetrating the stomata are indicated with an arrow. Calibration bars correspond to 25 μm.

The above qRT-PCR analyses do not resolve expression levels in individual cells. The reporter line harboring the mCherry gene expressed under the control of the *AvrStb6* promoter and located in the *AvrStb6* locus enabled us to monitor the *AvrStb6* expression pattern at the cellular level. Confocal microscopy pictures were taken at 6 dpi to maximize the number of penetration events of *Z. tritici*, while minimizing the autofluorescence produced by plants at later time points of the infection. The *AvrStb6* promoter shows little activity during hyphal growth on the plant surface but is strongly activated in hyphae approaching the stomata (Figure 6D), as previously demonstrated (10). We investigated whether this expression pattern was mediated by KATs. Remarkably, all the analyzed mutants grew as hyphae on the leaf surface and reached leaf stomata. Interestingly, at 6 dpi, Δ*Sas3* showed only minimal activation of the *AvrStb6* promoter even in hyphae attempting to penetrate the stomata (Figure 6F), which confirms the previous observation of Sas3 being involved in effector gene regulation during infection. On the other hand, the activity of the *AvrStb6* promoter in Δ*Elp3* was higher than in the control regardless of the proximity to stomata (Figure 6I). Δ*Sas2*, Δ*Ngs1* and Δ*Gcn5* displayed a similar *AvrStb6* expression pattern than the control at 6 dpi (Figures 6E, 6G and 6H). The results demonstrate that Sas3 is involved in effector gene upregulation during stomata penetration in *Z. tritici*.

### 2.8 Sas3 contributes to H3K9 and H3K14 acetylation of effector loci during plant infection

We subsequently evaluated whether Sas3-mediated expression regulation of effector genes is associated with histone acetylation during plant infection. We determined the acetylation levels of H3K9 and H3K14 during wheat infection in Δ*Sas3* lines. We observed a reduction in the relative acetylation levels of H3K9 (Figure 7A) and H3K14 (Figure 7B) in the *AvrStb6* promoter region and in *Avr3D1* in Δ*Sas3* compared to the control line during plant infection. As expected, acetylation levels of H3K9 and H3K14 in the control locus (*TFIIIC*) and 1000 bp upstream of the start codon of *AvrStb6* were not affected by the *Sas3* deletion. These results demonstrate that Sas3 is involved in plant-associated acetylation of H3K9 and H3K14 in effector loci. We suggest that the reduced effector transcript levels and the hindered infection of Δ*Sas3* (Figures 4 and 6) might be due to a reduction in histone acetylation (Figure 7).

**Figure 7.**
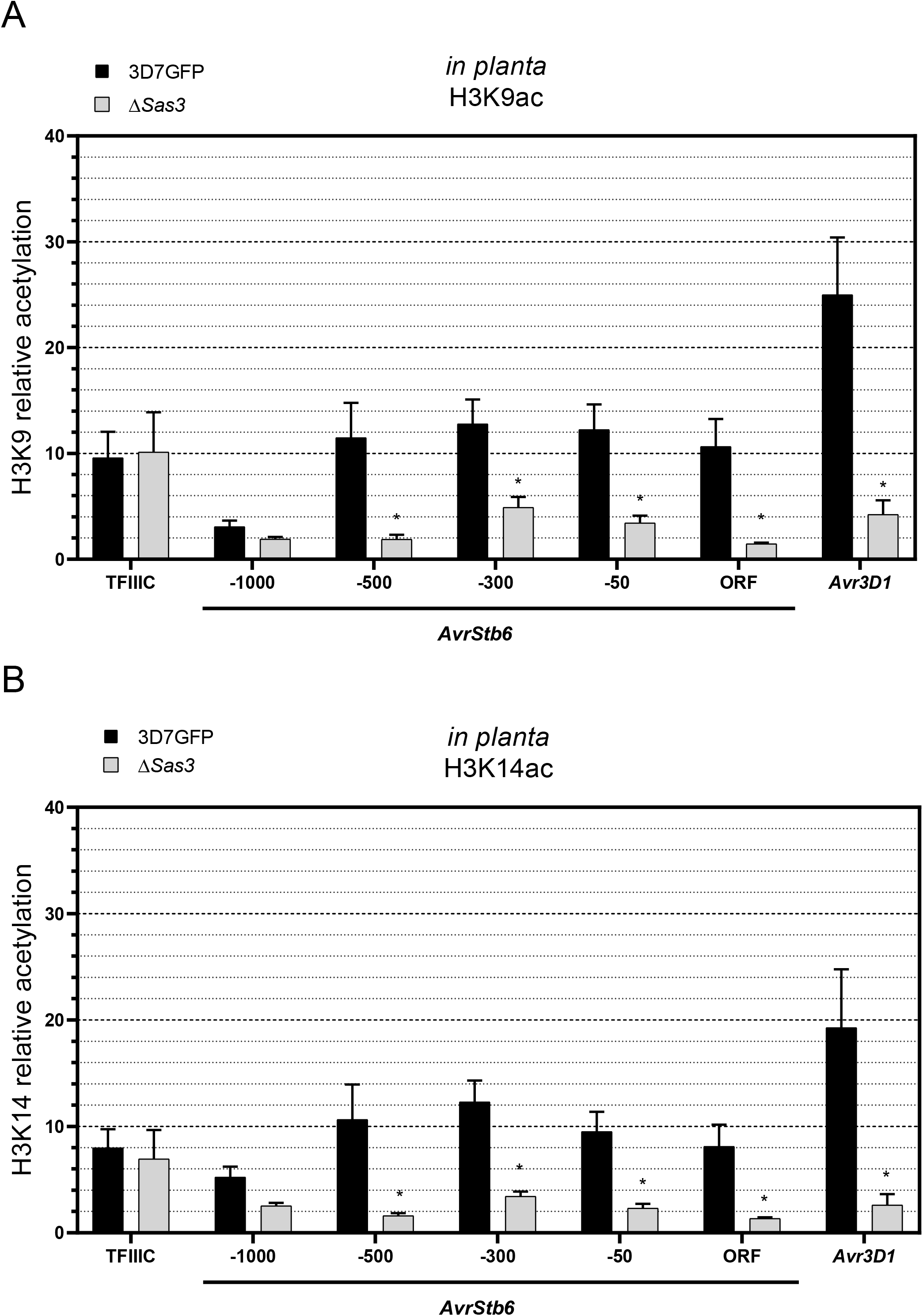
Histone H3 lysine 9 (H3K9) and 14 (H3K14) acetylation in effector genes is mediated by Sas3 *in planta*. Relative acetylation of H3K9 (A) and H3K14 (B) in the control (3D7-GFP) and Δ*Sas3* in different regions of *AvrStb6*: 1000 base pairs (bp; −1000), 500 base pairs (−500), 300 bp (−300) and 50 bp (−50) upstream the start codon; and in the open reading frame region (ORF). We also evaluated the acetylation of these two marks 300 bp upstream of the starting codon of *Avr3D1. TFIIIC* (100 bp upstream of start codon) was used as control. Chromatin immunoprecipitation experiments were performed during host colonization at 11 dpi. Acetylation levels are shown relative to H3 levels. Bars show the average of three independent biological replicates and the error bars represent the standard error of the mean. Asterisks indicate significant differences between Δ*Sas3* and 3D7-GFP according to two-way ANOVA and Bonferroni tests (p < 0.05).

## 3. DISCUSSION

Exploring the mechanisms by which plant pathogens activate their infection machinery is key for understanding how the interaction between the host and the pathogen is established. In the past years, chromatin remodeling has been shown to be crucial for effector gene activation during plant infection (10, 11, 13, 33). However, the specific chromatin modifications that are involved in this activation remain largely unknown. In this work we investigated the role of KATs and histone acetylation in the virulence of the fungal plant pathogen *Z. tritici*. We demonstrated that Sas3-mediated histone acetylation dynamics are associated with upregulation of effector genes during plant infection.

A total of three KATs from the MYST family were identified in *Z. tritici*, Sas3, Sas2 and Esa1. We showed that Sas3 and Sas2 are involved in the expression regulation of well-characterized effector genes, such as *AvrStb6*. Although Sas2 did not affect asexual reproduction and the speed at which necrotic lesions developed, it clearly shaped the visual appearance of lesions, manifested by red-orange spots in lesions produced in infections by the Δ*Sas2* mutant. We considered that this altered symptom development might be the result of misexpression of *AvrStb6* and potentially other effector genes in the Δ*Sas2* mutant. Likewise, the Sas2 orthologue in *B. cinerea* (BcSas2) is involved in regulation of virulence (34), suggesting a conserved role of Sas2 in effector gene activation in fungal pathogens. Disruption of *Sas3* in *Z. tritici* led to a reduction of virulence and pycnidia formation. Similarly, Sas3 from *M. oryzae* is involved in virulence (18). We additionally noticed that Sas3 is required for normal growth under axenic conditions since colony size was reduced in Δ*Sas3*. Although this reduction in growth could indirectly lead to a reduction of virulence, we suggest that Sas3 is directly involved in the regulation of virulence since (i) Δ*Sas3* grows as hyphae on the leaf surface of wheat and are able to reach the stomata, (ii) *AvrStb6* activation is impaired in the proximity to the stomata in Δ*Sas3* mutants, and (iii) misactivation of effector genes occurs at stages of infection when the fungal biomass is similar to the control (6 dpi). Thus, we believe that the impaired virulence of Δ*Sas3* mutants is most likely a consequence of effector gene misregulation, featuring reduced expression levels of *AvrStb6, AvrStb9*, and *Mycgr3G76589* during plant infection, highlighting the contribution of Sas3 in the activation of effector genes. (30). Remarkably, mutants in Sas3 are impaired in histone acetylation of effector genes during plant infection, suggesting that Sas3-mediated acetylation is a crucial mechanism driving the transcriptional reprogramming of effector genes during plant infection *Z. tritici* (16).

We identified three members of the GNAT KAT family in *Z. tritici*, with distinct roles in the infection cycle. Expression analysis of effector genes in GNAT mutants revealed that GNATs might be involved in negatively regulating effector genes. However, GNATs might not directly regulate effector genes but rather other transcriptional regulators. We suggest that in Δ*Gcn5* the higher expression levels of effector genes might lead to faster development of necrosis. This might be due to two possible scenarios. The high accumulation of effectors might lead to an early recognition of the pathogen by the host, resulting in a strong immune response which might be manifested by cell death. Alternatively, misregulation of cell-death inducing effectors might directly produce necrosis at earlier stages of the infection. Additionally, we showed that Gcn5 is involved in asexual reproduction since the knockout mutant developed relatively few pycnidia. Interestingly, Gcn5 negatively regulates symptom development but positively regulates reproduction, supporting that different mechanisms govern virulence and pycnidia production, as previously shown (23).

H3K9 and H3K14 acetylation are well-known euchromatic marks (35). Accordingly, we observed an increase in histone acetylation levels in *Avr3D1* and *AvrStb6* during plant infection, along with the derepression of these two effector genes. We suggest that this increase in histone acetylation levels contributes to the *in planta*-specific upregulation of effector genes in *Z. tritici*. Previously, we reported a decrease in the levels of H3K27 and H3K9 trimethylation in effector loci during plant infection associated with effector gene depression (10). We consider that this reduction of histone repressive marks and an increase in activating marks, such as those described in the current work, promote a local switch from repressive to permissive chromatin, allowing the access of nucleosome remodeling complexes and structural modifications in chromatin, including decondensation of the chromatin fiber. Previous work demonstrated that for selected effector genes, such changes in chromatin structure are very local and do not affect neighboring loci (10), suggesting the targeted location of KATs to effector loci during plant infection. Substrate specificities have been reported to be mediated by certain subunits from KAT complexes or KAT domains that interact with nucleosomes (36). In *Z. tritici*, a bromodomain and a PHD-finger domain were identified in Gcn5 and Sas3, respectively, both with a potential role in substrate specificity or in interaction with regulatory proteins (37). In addition, the concerted expression of effector genes during plant infection most likely requires altered transcription factor activities or levels, as demonstrated for the Zn2Cys6 family member transcription factor Pf2 from *L. maculans*. In this case, the coordinated action of trimethylation of H3K9 and Pf2 governs the specific expression pattern of effector genes (38). Accordingly, we propose that chromatin modifications and still unknown transcription factors might jointly act as derepressors of effector genes of *Z. tritici* during plant infection.

We have shown that histone modifications, involving acetylation and demethylation (10), mediate the activation of effector genes during plant infection. Elucidating the crosstalk between histone modifications, their direct or indirect function in effector gene regulation and the role of classic transcriptional activators and repressors will help us to further understand the molecular mechanisms linking chromatin and stage-specific transcriptional changes. Future work aiming to unveil global changes in histone acetylation and methylation patterns during plant infection will shed more light on the contribution of these histone marks to the regulation of the infection machinery.

## 4. MATERIALS AND METHODS

### 4.1 Fungal and bacterial strains used

We used the *Z. tritici* Swiss strain ST99CH_3D7 ((39); abbreviated as 3D7). All mutants were obtained either in 3D7 expressing the codon optimized version of the enhanced green fluorescent protein (eGFP) (3D7-GFP; (10, 40)), or in a mutant reporter line that expresses *mCherry* fused to His1 under the control of the *AvrStb6* promoter and located within the *AvrStb6* locus (10). Stellar *Escherichia coli* HST08 cells (Takara Bio, Japan) and the *Agrobacterium tumefaciens* strain AGL1 were used for cloning and *Z. tritici* transformation, respectively.

### 4.2 Bioinformatic tools

To identify and classify the KAT orthologues from *Z. tritici*, we first used the basic local alignment search tool (BLAST; (41)) from the National Centre for Biotechnology Information (NCBI) using the previously characterized KAT protein sequences from *S. cerevisiae* as queries (Table S1). Reverse BLAST was also performed to confirm that the identified protein sequences in *Z. tritici* were KAT orthologues. In parallel, we used the dbHiMo web-based data browser (26). Results obtained in the BLAST analysis were compared with results using dbHiMo to confirm that the same KATs were found in both cases. We obtained a multiple sequence alignment of the KAT orthologues (MUSCLE; (42)) and a phylogenetic tree. The protein sequences used for constructing the phylogenetic tree were obtained from NCBI (Table S2). Phylogenetic trees were constructed using the Molecular Evolutionary Genetics Analysis (MEGA-X 10.2; (43)) software, applying the maximum likelihood (ML) method with 1000 non-parametric bootstraps as statistical support. Trees were rooted using the midpoint rooting method. We edited the trees using the Interactive Tree Of Life (iTOL; (44)) software. Additionally, we identified protein domains in *Z. tritici* HMEs using HMMER (45) including all databases (Pfam, TIGRFAM, Gene3D, Superfamily, PIRSF and TreeFam) and represented the different protein domains using the R package “ragp” (46).

### 4.3 Generation of *Z. tritici* transformants

Plasmids for targeted gene deletion by homologous recombination were assembled using the In-Fusion^®^ HD Cloning Kit (Takara Bio, Japan). Briefly, the nourseothricin resistance gene PCR-amplified from pES1-NAT-GFP (47) was flanked by homology arms of ca. 1 kb and inserted into the KpnI-SbfI-linearized acceptor plasmid pCGEN (48). Similarly, constructs for genetic complementation were generated by assembling gene sequences spanning from ca. 1 kb upstream of the start codon to right before the stop codon and the C-terminal 4xMyc-tag (49) into XhoI-linearized pLM1. Primers are listed in Table S3. *Z. tritici* gene deletion mutants and respective complementation mutants were obtained by *A. tumefaciens*-mediated transformation as previously described (10, 50) using nourseothricin (25 μg·mL^−1^) and hygromycin (100 μg·mL^−1^) for selection, respectively. Despite the presence of homology arms in the T-DNA, transformation of *Z. tritici* typically yields high frequencies of ectopic insertions by non-homologous end-joining instead of or in addition to targeted insertions by homologous recombination. To distinguish deletion mutants from ectopic insertion mutants, a PCR-based mutant screening was performed using either purified genomic DNA as template or directly adding liquid culture to the PCR reaction. The screening method includes a primer binding site present between the nourseothricin resistance gene and the downstream homology arm. The sequence of this screening primer binding site was chosen to match the sequence of the gene to be deleted in a way that yields two distinct amplicons in deletion and ectopic insertion mutants when combined with a primer binding site located in the region downstream of the homology arm (Figure S9). Since this screening method yields distinct amplicons for both deletion and ectopic insertion mutants, failed PCR reactions can easily be identified by the lack of both amplicons. Furthermore, the presence of both amplicons in the same reaction allows the identification of impure mutant lines and heterokaryons. Insertion copy numbers were determined by qPCR and mutant lines with multiple inserts were discarded. At least two independent lines were obtained per mutant and used for subsequent experiments.

### 4.4 Infection assays

Infection assays were performed on wheat (*Triticum aestivum* L.) plants of cultivar Runal grown for 15 days at 18°C during the day and 15°C during the night, with 16 hours of light and 65% of relative humidity. Sixteen seeds of cultivar Runal were sown in 11×11×12 cm pots with a peat-based substrate. Plants were fertilized after one week (Universal fertilizer, COMPO, Münster, Germany). The fungal inoculum was prepared one week before the infection by inoculating 50-100 μL of glycerol stock in 50 mL of on yeast extract-peptone-dextrose broth (YPD; yeast extract 10 g·L^−1^; peptone 20 g·L^−1^; dextrose 20 g·L^−1^) amended with kanamycin (50 μg·mL^−1^). Spore suspensions were prepared as described (24) and quantified using a BLAUBRAND bright-line Neubauer improved hemocytometer (0.100 mm depth, 0.0025 mm^2^ area; Brand, Wertheim, Germany), except for the ChIP-qPCR experiment in which we used the Spore Counter macro vs 2.13 (https://github.com/jalassim/SporeCounter.git; Julien Alassimone; ETH-Zürich, Switzerland).

Wheat infection assays were performed using fungal suspensions at a concentration of 10^7^ spores·mL^−1^ in 0.1% Tween-20 as previously described (24). Each pot was sprayed either with 12.5 mL 0.1% Tween-20 for mock treatment or 12.5 mL spore suspension for controls and mutants. At least two independent mutant lines were used to evaluate symptom development and pycnidia production. Symptoms produced by *Z. tritici* were analyzed on the second leaf at two different time point using ImageJ (51) and an automated image analysis method (52). The percentage of leaf aPLACL and pycnidia counts per square centimeter of lesion were used as proxy for virulence and asexual reproduction, respectively.

### 4.5 Developmental assays

We performed fitness assays with *Z. tritici* mutants. A 3-μL drop of *Z. tritici* spore suspensions at 10^6^, 10^5^, 10^4^ and 10^3^ spores·mL^−1^ was placed on different types of media: YMA, YMA supplemented with NaCl (0.5 M), H_2_O_2_ (1 mM), sorbitol (1 M), calcofluor white (200 ng·μL^−1^), or Congo red (2 mg·mL^−1^); minimal medium (MM; Voguel’s medium (53)), and MM supplemented with fructose (5 g·L^−1^), galactose (50 mM), GlcNAc (2.5 mM), or glucose (2.5 mM). Inoculated agar plates were incubated at 18°C. An additional plate of YMA was incubated at 28°C. Pictures were taken after 6 days.

Area of individual colonies of the mutants was estimated by inoculating ca. 100 colony-forming units on YMA. Three independent replicates of each mutant were performed. After 5 days of incubation at 18°C, pictures of the plates were taken, and the colony size was analyzed using ImageJ.

### 4.6 Confocal laser scanning microscopy assays

Confocal assays were performed on a Zeiss LSM 880 super-resolution confocal microscope with fast Airyscan. The emission settings were: 511 to 564 nm for the eGFP channel, 603 to 623 nm for the mCherry channel, 460 to 480 nm for the mTurquoise2 channel and 692 to 697 nm for the chloroplast detection. For excitation, an argon (488 nm) laser was used for track 1 (mCherry, and chloroplasts) and a diode laser (405 nm) was used for track 2 (mTurquiose2 and eGFP). Image processing was performed using Fiji (54) and included generation of maximum-intensity Z-projections for merging channels and addition of calibration bars. Colors were selected manually to facilitate channel identification. At least two independent lines per transformant were used. Experiments were performed at least twice.

### 4.7 Effector gene expression analysis

Axenically grown cultures obtained as described above were filtered through a nylon membrane and centrifuged at 5000 g, 4°C for 5 minutes. Supernatant was discarded and fungal pellets were ground in liquid N2, using mortar and pestle. Infected plant tissue was collected at 6- and 10-days post infection (dpi). Eight centimeters of second leaves (after discarding 2 cm from the tip) were used for RNA extraction. Each replicate consisted of 2 leaves. At least three biological replicates were analyzed per treatment. RNA was extracted with Trizol (Life Technologies), purified (RNAeasy Mini Kit, QIAGEN Inc., The Netherlands) and treated with DNAse (QIAGEN Inc., The Netherlands). cDNA was synthesized by using the Primescript RT reagent kit (Takara Bio, Japan). qPCR was performed in a LightCycler480 II (Roche Diagnostics International AG, Rotkreuz, Switzerland) using the primers listed in Table S3, and data were analyzed with the LightCycler 480 software (Roche Diagnostics International AG, Rotkreuz, Switzerland) using *histone H3 (3D7.g6784*) and *beta tubulin (3D7.g2064*) as reference genes. Relative fungal biomass *in planta* was calculated by dividing the transcript values of fungal housekeeping genes (histone H3 and beta tubulin) and the plant housekeeping gene *TaCDC48 (Triticum aestivum* cell division control protein 48 homolog E-like; *Traes4A02G035500*) (55).

### 4.8 Chromatin extraction and immunoprecipitation (ChIP)

For *in vitro* chromatin extraction 150 mg of tissue were used, while for *in planta* chromatin extraction 250 mg of tissue were used. The micrococcal nuclease (M0247S; New England Biolabs, Ipswich, MA, USA) reaction was performed at 37°C for 20 minutes. Chromatin fixation, immunoprecipitation and de-crosslinking were performed as previously described (9, 10). Antibodies anti-H3 (ab1791, Abcam, Cambridge, UK), anti-H3K9ac (ab10812, Abcam, Cambridge, UK) and anti-H3K14ac (ab52946, Abcam, Cambridge, UK) were applied in 1:200 ratio. Subsequent qPCR was carried out on a LightCycler 480 instrument (Roche Diagnostics International AG, Rotkreuz, Switzerland). Acetylation levels were estimated as relative levels of H3K9ac and H3K14ac normalized to histone H3 as previously described (56, 57).

### 4.9 Statistics

Statistical analysis and graphic representations were performed using either RStudio version 1.4.1717 (58) or GraphPad Prism 8.0.2 for Windows (GraphPad Software, San Diego, California). For conducting the statistical analyses, Gaussian distribution of the data was tested using a Shapiro-Wilk normality test and homogeneity of variances was analyzed using a Brown-Forsythe test. If the data followed a normal distribution and preserved homoscedasticity, the parametric ordinary one-way ANOVA test was applied together with Fisher’s LSD test (p-value < 0.05). If the aforementioned assumptions were not met, the non-parametric Kruskal-Wallis test was applied together with Dunn’s uncorrected test (p-value < 0.05). In the case of ChIP data, two-way ANOVA and Bonferroni analyses were performed (p-value < 0.05). All raw data used for performing main text and supplementary figures are available in Dataset S1 and S2, respectively.

## 5. ACKNOWLEDGEMENTS

We would like to specially thank Javier Barrero, Pedro Crevillen and Jose Antonio Abelenda for their support and help with ChIP experiments and results interpretation. We would like to thank Thierry Marcel and Reda Amezrou for providing us with nonpublished information and to Julien Alassimone for providing us the Spore Counter macro. We thank Gero Steinberg for providing us with the vector containing the codon-optimized eGFP and 3D7-GFP, Jason Rudd for providing the vector PCGEN and Manuel Ene Ordorica for providing the 4xmyc sequence. We thank the Zymoseptoria laboratory at the CBGP for their help and support. We thank DSP Ltd (Delley, Switzerland) for providing us wheat seeds. The research was financed by the Ministry of Science and Innovation (Grant PID2019-108693RA-I00 financed by MICIN/AEI/ to AS-V). ASV was recipient of Ramon y Cajal grant RYC2018-025530-I of Spanish Ministry of Science, Innovation and Universities (MCIN/AEI/ and El FSE). MSF was recipient of Margarita Salas financed by the European Union - Next Generation EU (Grant MARSALAS21-31).

## 6. DATA AVALAIBILITY STATEMENT

The authors declare that the raw data of all the experiments are included in supplemental material (Datasets S1 and S2).

## 9. SUPPLEMENTAL MATERIAL TABLES

**Table S1.** BLASTp analysis identify 5 lysine acetyltransferases (KATs) from the MYST and GNAT families in *Z. tritici*. KATs of *Saccharomyces cerevisiae* used, query cover, E-value, percentage of identity, accession number and gene ID of the best hit in *Z. tritici* are indicated.

| *S. cerevisiae*<br>KAT | Query<br>Cover | E- value | Percentage<br>of Identity | Accession<br>(*Z. tritici*) | Associated *Z. tritici*<br>gene ID |
| --- | --- | --- | --- | --- | --- |
| Gcn5 | 80% | 2e-164 | 60.76% | SMQ49624.1 | 3D7.g4775 |
| Elp3 | 99% | 0.0 | 74.24% | SMQ53347.1 | 3D7.g8500 |
| Sas2 | 60% | 1e-46 | 39.71% | SMQ51878.1 | 3D7.g7031 |
| Sas3 | 47% | 9e-79 | 36.83% | SMQ49112.1 | 3D7.4263 |
| Esa1 | 99% | 5e-162 | 49.41% | SMQ54127.1 | 3D7.9281 |

**Table S2.** NCBI accession numbers used for performing the phylogenetic tree.

| **KAT** | **Organism** | **NCBI accession number** |
| --- | --- | --- |
| **Ngs1** | *Neurospora crassa* | CAE85548.1 |
| **Ngs1** | *Fusarium oxysporum* | KAF6521047.1 |
| **Ngs1** | *Trichoderma reesei* | XP_006966911.1 |
| **Ngs1** | *Aspergillus fumigatus* | KAH3100114.1 |
| **Ngs1** | *Histoplasma capsulatum* | QSS62698.1 |
| **Ngs1** | *Zymoseptoria tritici* | SMQ47703 |
| **Ngs1** | *Candida albicans* | AOW30790.1 |
| **Sas2** | *Saccharomyces cerevisiae* | DAA10024.1 |
| **Sas2** | *Fusarium oxysporum* | SCO82216.1 |
| **Sas2** | *Kluyveromyces marxianus* | XP_022675924 |
| **Sas2** | *Beauveria bassiana* | KAF1731046.1 |
| **Sas2** | *Zygosaccharomyces mellis* | GCE98240.1 |
| **Sas2** | *Fusarium graminearum* | XP_011324667.1 |
| **Sas2** | *Zymoseptoria tritici* | SMQ51878.1 |
| **Sas3** | *Saccharomyces cerevisiae* | DAA07067.1 |
| **Sas3** | *Fusarium graminearum* | XP_011320283.1 |
| **Sas3** | *Metarhizium robertsii* | XP_007818471.1 |
| **Sas3** | *Fusarium oxysporum* | EWZ39814.1 |
| **Sas3** | *Magnaphorte oryzae* | XP_003713627.1 |
| **Sas3** | *Zymoseptoria tritici* | SMQ49112.1 |
| **Esa1** | *Saccharomyces cerevisiae* | DAA11012.1 |
| **Esa1** | *Magnaphorte oryzae* | XP_003719696.1 |
| **Esa1** | *Fusarium graminearum* | Q4IEV4.1 |
| **Esa1** | *Neurospora crassa* | XP_962217.1 |
| **Esa1** | *Zymoseptoria tritici* | SMQ54127.1 |
| **Gcn5** | *Saccharomyces cerevisiae* | DAA07067.1 |
| **Gcn5** | *Ustilago maydis* | CAC80426.1 |
| **Gcn5** | *Fusarium oxysporum* | EWZ52160.1 |
| **Gcn5** | *Neurospora crassa* | XP_001728480.2 |
| **Gcn5** | *Schizosaccharomyces pombe* | Q9UUK2.1 |
| **Gcn5** | *Zymoseptoria tritici* | SMQ49624.1 |
| **Elp3** | *Saccharomyces cerevisiae* | DAA11347.1 |
| **Elp3** | *Fusarium graminearum* | XP_011317913.1 |
| **Elp3** | *Magnaphorte oryzae* | XP_003710346.1 |
| **Elp3** | *Schizosaccharomyces pombe* | NP_594862.1 |
| **Elp3** | *Fusarium oxysporum* | XP_018239078.1 |
| **Elp3** | *Candida albicans* | KAF6072097.1 |
| **Elp3** | *Zymoseptoria tritici* | SMQ53347.1 |

**Table S3.**
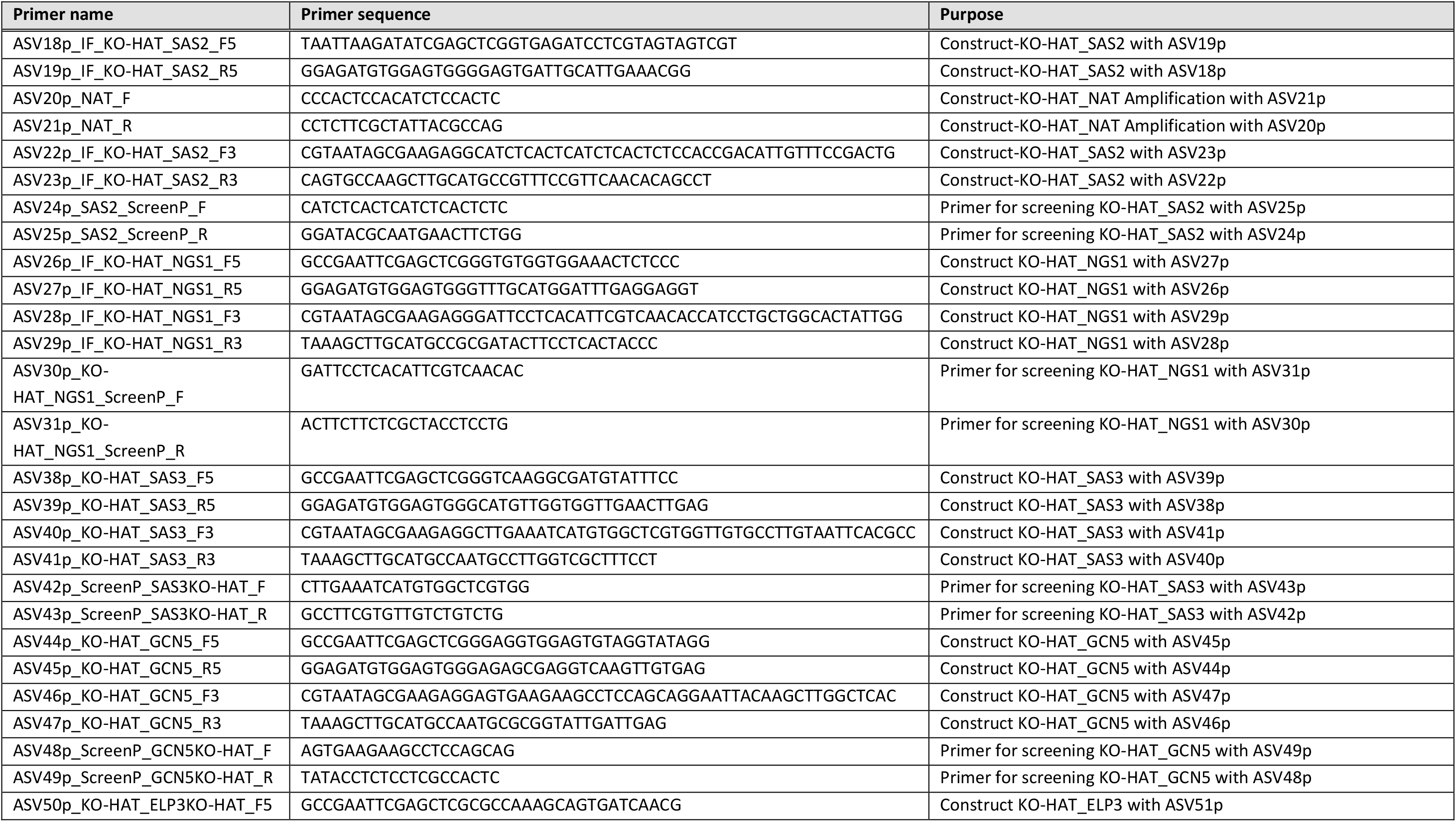
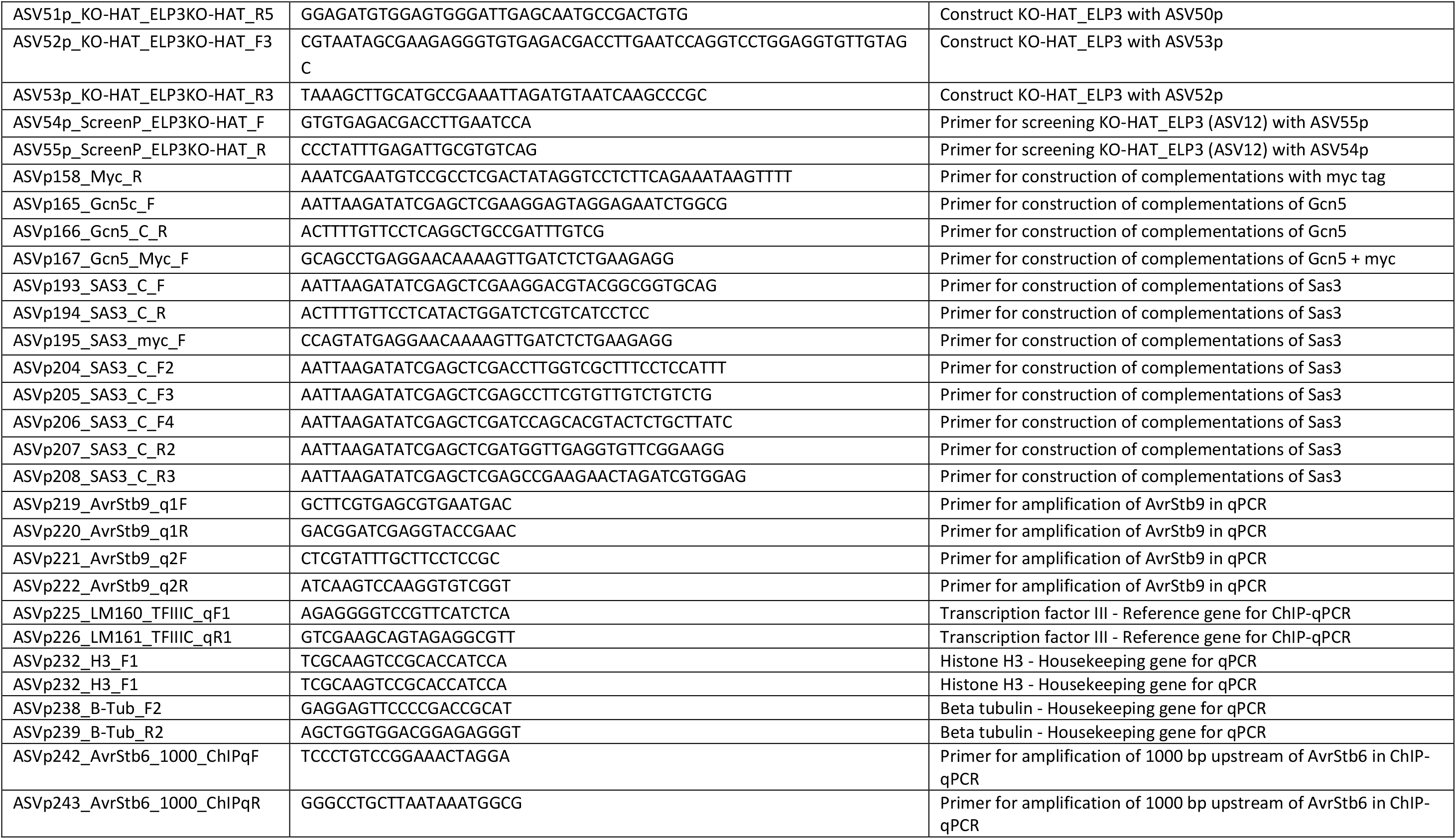
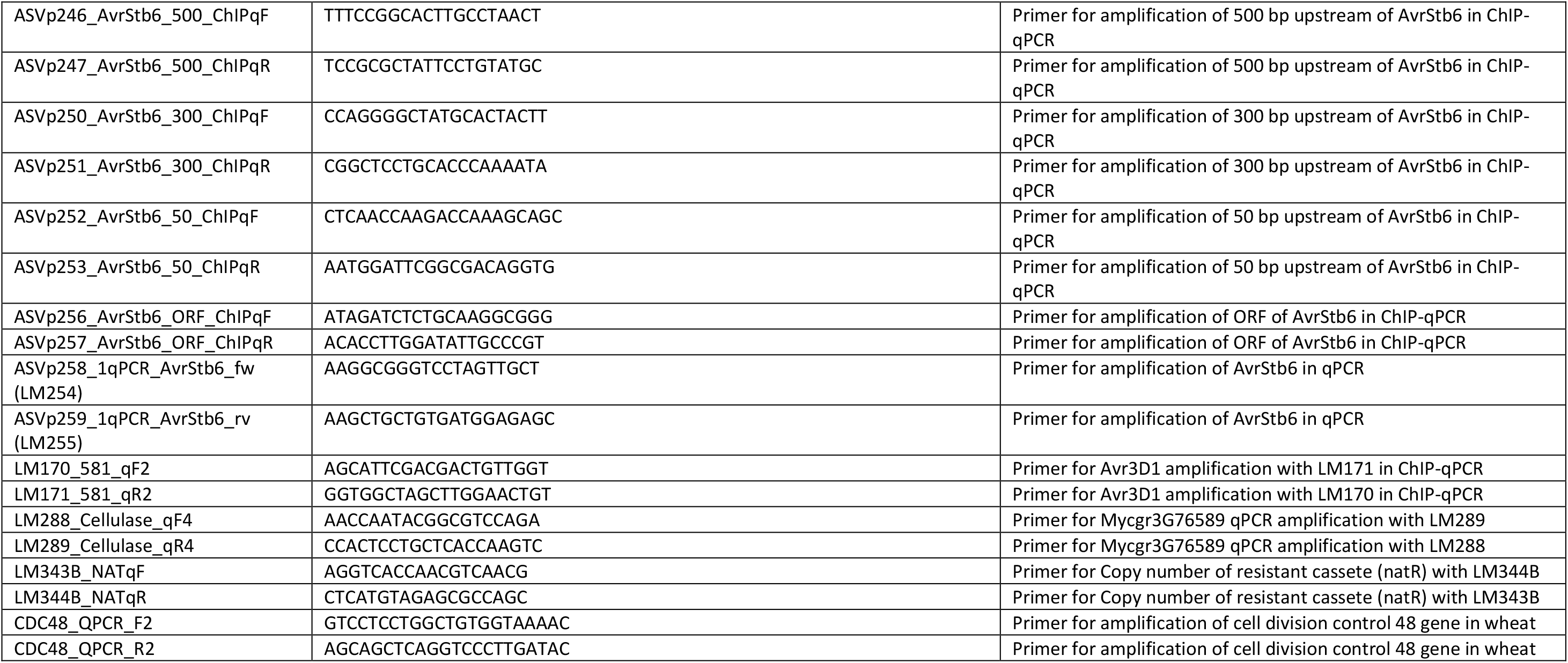
Primers used in this work.

## 10. SUPPLEMENTAL MATERIAL FIGURE LEGENDS

**Figure S1.**
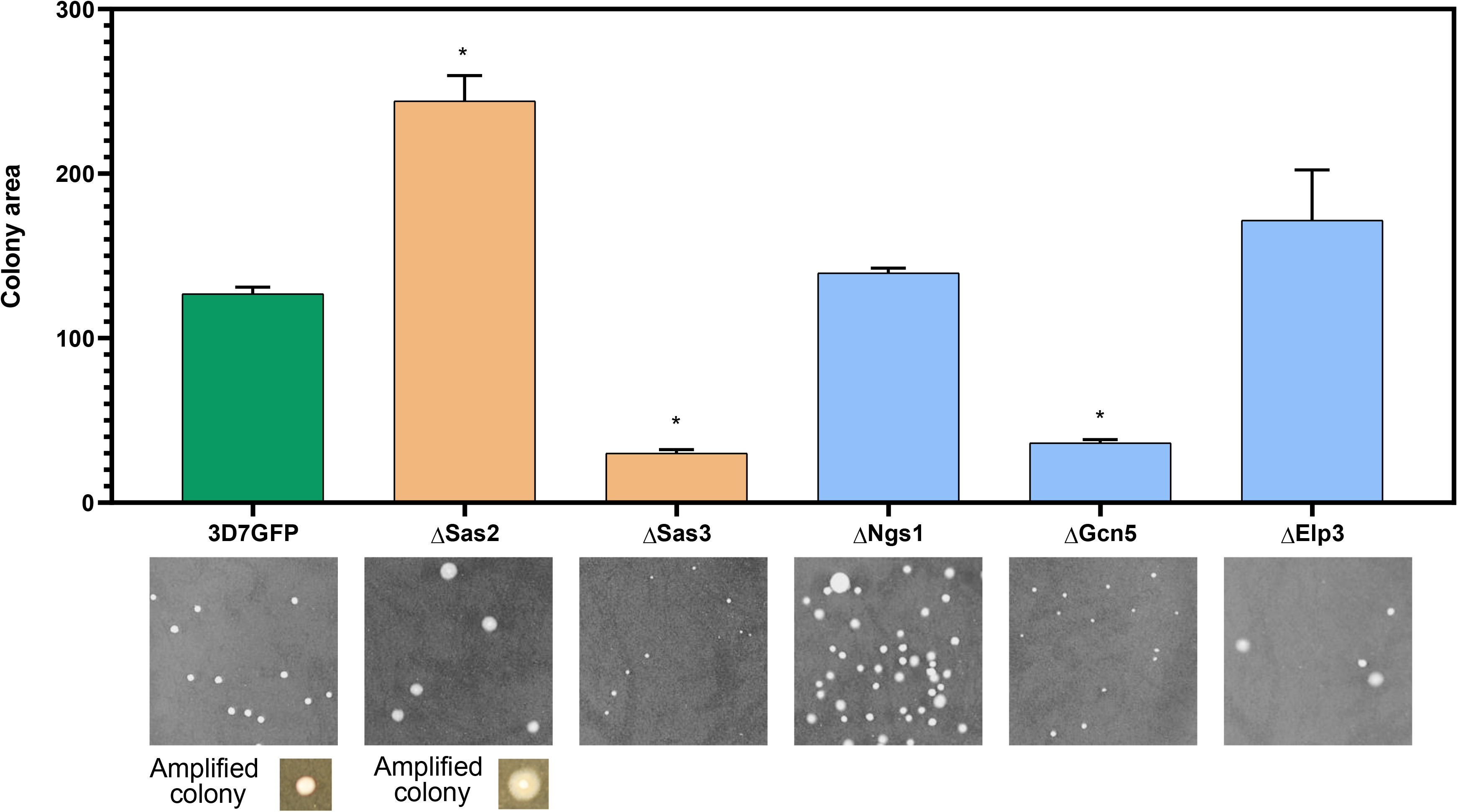
Sas3, Sas2 and Gcn5 regulate growth and development of *Z. tritici*. Colony area of the control (3D7-GFP), Δ*Ngs1*, Δ*Sas2*, Δ*Sas3*, Δ*Gcn5* and Δ*Elp3* grown for 5 days on yeast-malt-sucrose agar (YMA). Green bar represents 3D7-GFP, orange bars represent MYST-family mutants and blue bars represent GNAT-family mutants. Bars represent average of three independent biological replicates and error bars represent the standard error of the mean. A representative image of colonies of each mutant is shown. Additionally, an amplification of a colony of the control and Δ*Sas2* are shown. Asterisks indicate significant differences with 3D7-GFP according to Kruskal-Wallis and uncorrected Dunn’s tests (p < 0.05).

**Figure S2.**
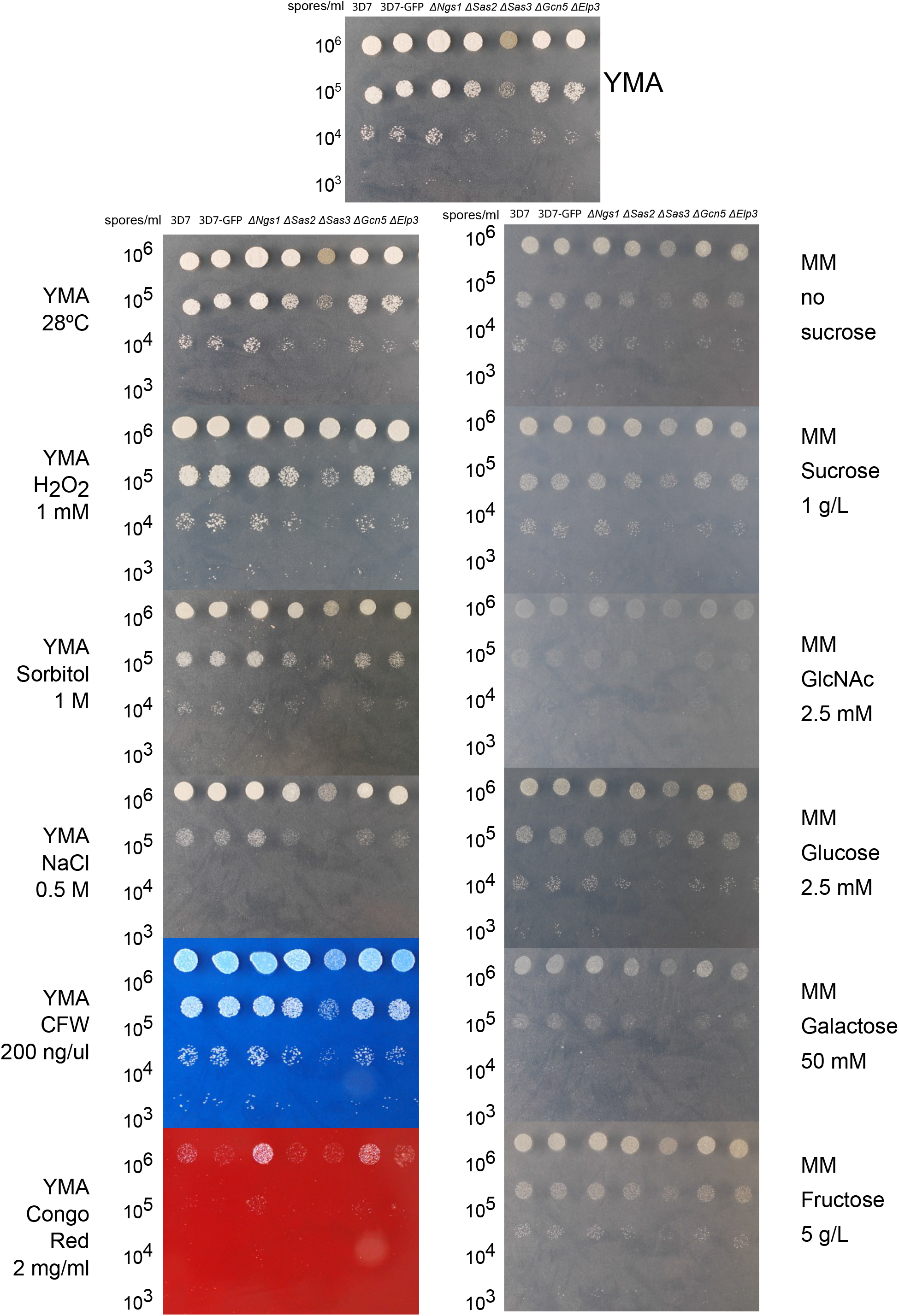
Lysine acetyltransferases (KATs) are not positive regulators of stress tolerance. Three μL of fungal spore suspensions at a concentration of 10^6^, 10^5^, 10^4^ and 10^3^ spores·mL^−1^ of the controls (3D7 and 3D7-GFP), Δ*Ngs1*, Δ*Sas2*, Δ*Sas3*, Δ*Gcn5* and Δ*Elp3* were inoculated. Media used were yeast-malt-sucrose agar (YMA); YMA supplemented with NaCl (0.5 M), H_2_O_2_ (1 mM), sorbitol (1 M), Calcofluor white (200 ng·μL^−1^) or Congo red (2 mg·mL^−1^); minimal medium (MM; Voguel’s), MM supplemented with fructose (5 g·L^−1^), galactose (50 mM), GlcNAc (2.5 mM), or glucose (2.5 mM). Plates were incubated at 18°C for 6 days. One additional plate of YMA was incubated at 28°C.

**Figure S3.**
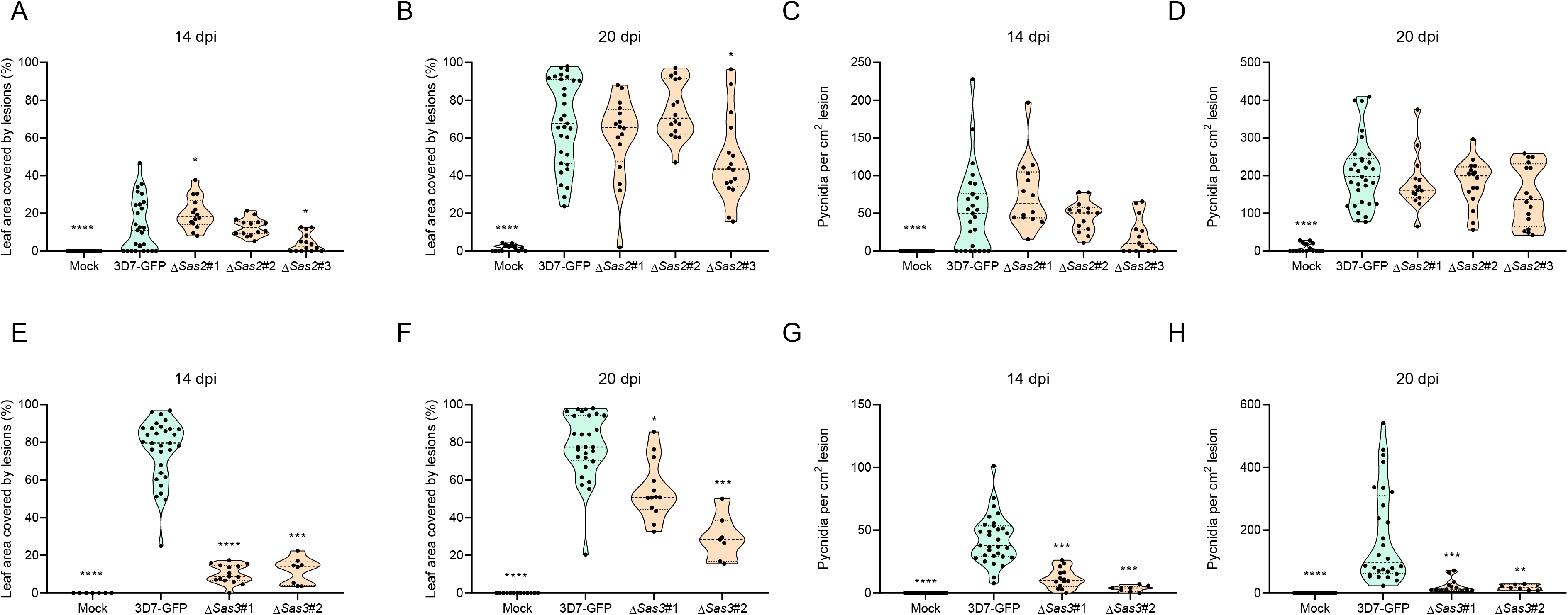
Infection assays of at least two independent mutant lines of lysine acetyltransferases (KATs) from the MYST family. Percentage of leaf area covered by lesions (PLACL) at 14 days post infection (dpi) (A) and at 20 dpi (B) and pycnidia per cm^2^ lesion at 14 dpi (C) and 20 dpi (D) in the three independent Δ*Sas2* lines (#1, #2, #3). PLACL at 14 dpi (E) and 20 dpi (F) and pycnidia per cm^2^ lesion at 14 dpi (G) and at 20 dpi (H) of two independent Δ*Sas3* mutant lines (#1, #2). Dashed lines represent the median, dotted lines represent first and third quartiles and black dots represent individual data points. Asterisks indicate statistically significant differences with the control (3D7-GFP) according to Kruskal-Wallis non-parametric statistical and posthoc uncorrected Dunn’s tests (* p < 0.05; ** p < 0.01; *** p < 0.001; **** p < 0.0001).

**Figure S4.**
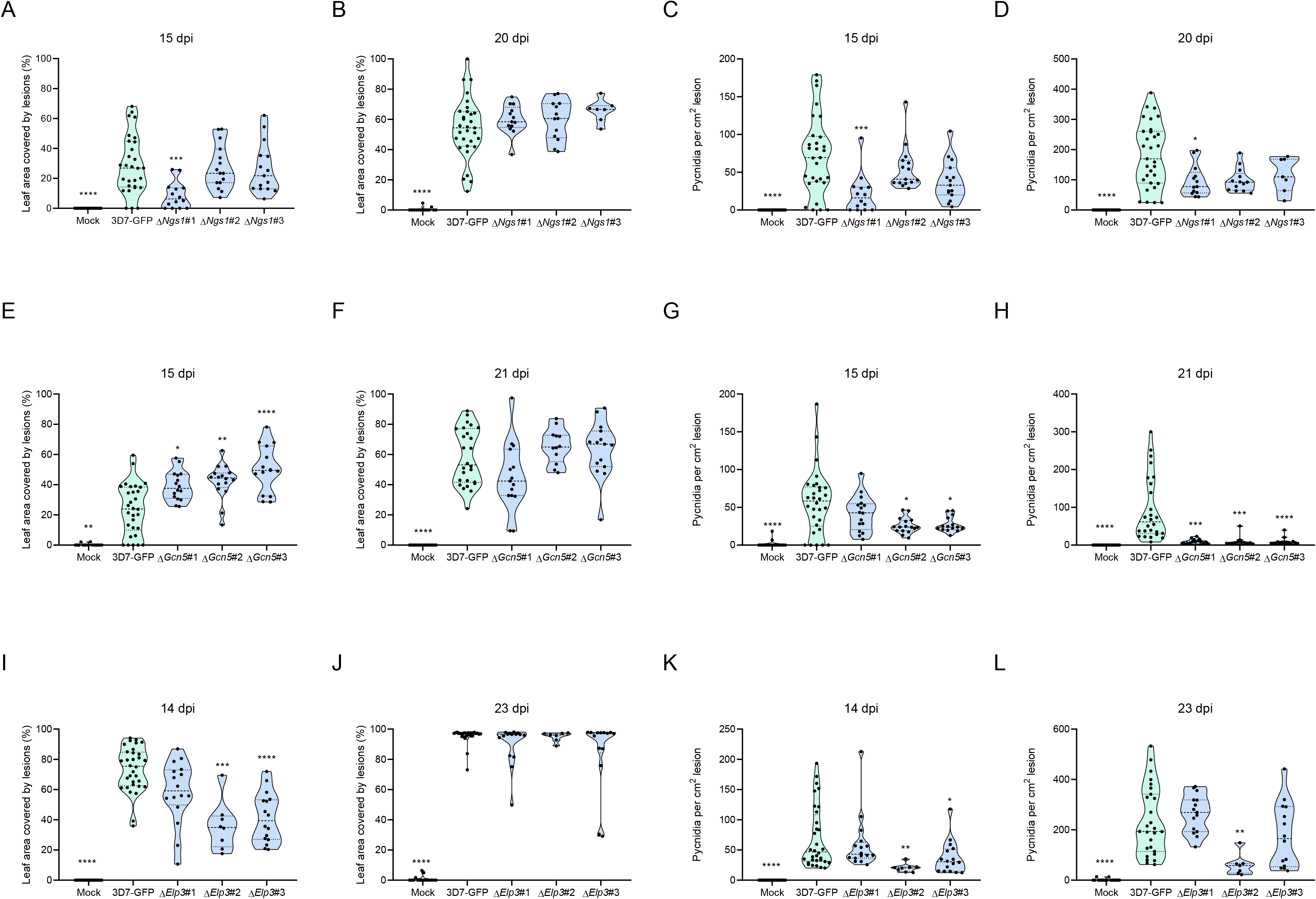
Infection assays of three independent mutant lines of lysine acetyltransferases (KATs) from the GNAT family. Percentage of leaf area covered by lesions (PLACL) at 15 days post infection (dpi) (A) and at 20 dpi (B) and pycnidia per cm^2^ lesion at 15 dpi (C) and 20 dpi (D) in three Δ*Ngs1* independent lines (#1, #2, #3). PLACL at 15 dpi (E) and 21 dpi (F) and pycnidia per cm^2^ lesion at 15 dpi (G) and at 21 dpi (H) of three independent lines of Δ*Gcn5* (#1, #2, #3). PLACL at 14 dpi (I) and 23 dpi (J) and pycnidia per cm^2^ lesion at 14 dpi (K) and at 23 dpi (L) of three independent lines of Δ*Elp3* (#1, #2, #3). Dashed lines represent the median, dotted lines represent first and third quartiles and black dots represent individual data points. Asterisks indicate statistically significant differences with 3D7-GFP according to the Kruskal-Wallis non-parametric statistical and posthoc uncorrected Dunn’s tests (* p < 0.05; ** p < 0.01; *** p < 0.001; **** p < 0.0001).

**Figure S5.**
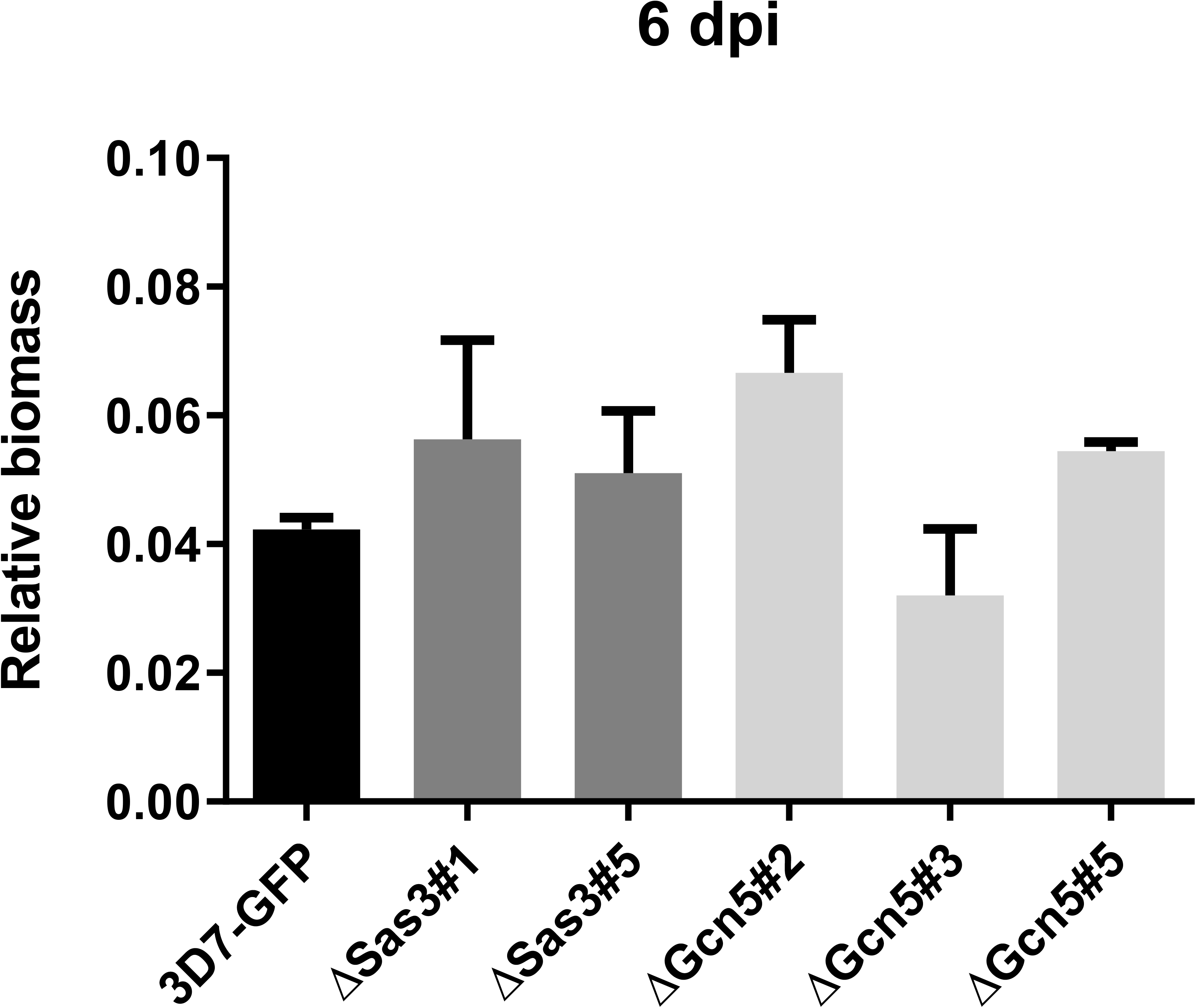
Fungal biomass of Δ*Sas3* and Δ*Gcn5*. Relative fungal biomass of the control (3D7-GFP), two independent lines of Δ*Sas3* and three independent lines of Δ*Gcn5* at 6 days post infection (dpi). Bars correspond to the average of three biological replicates. Error bars represent the standard error of the mean. No significant differences with 3D7-GFP according to the Kruskal-Wallis test were identified (p < 0.05).

**Figure S6.**
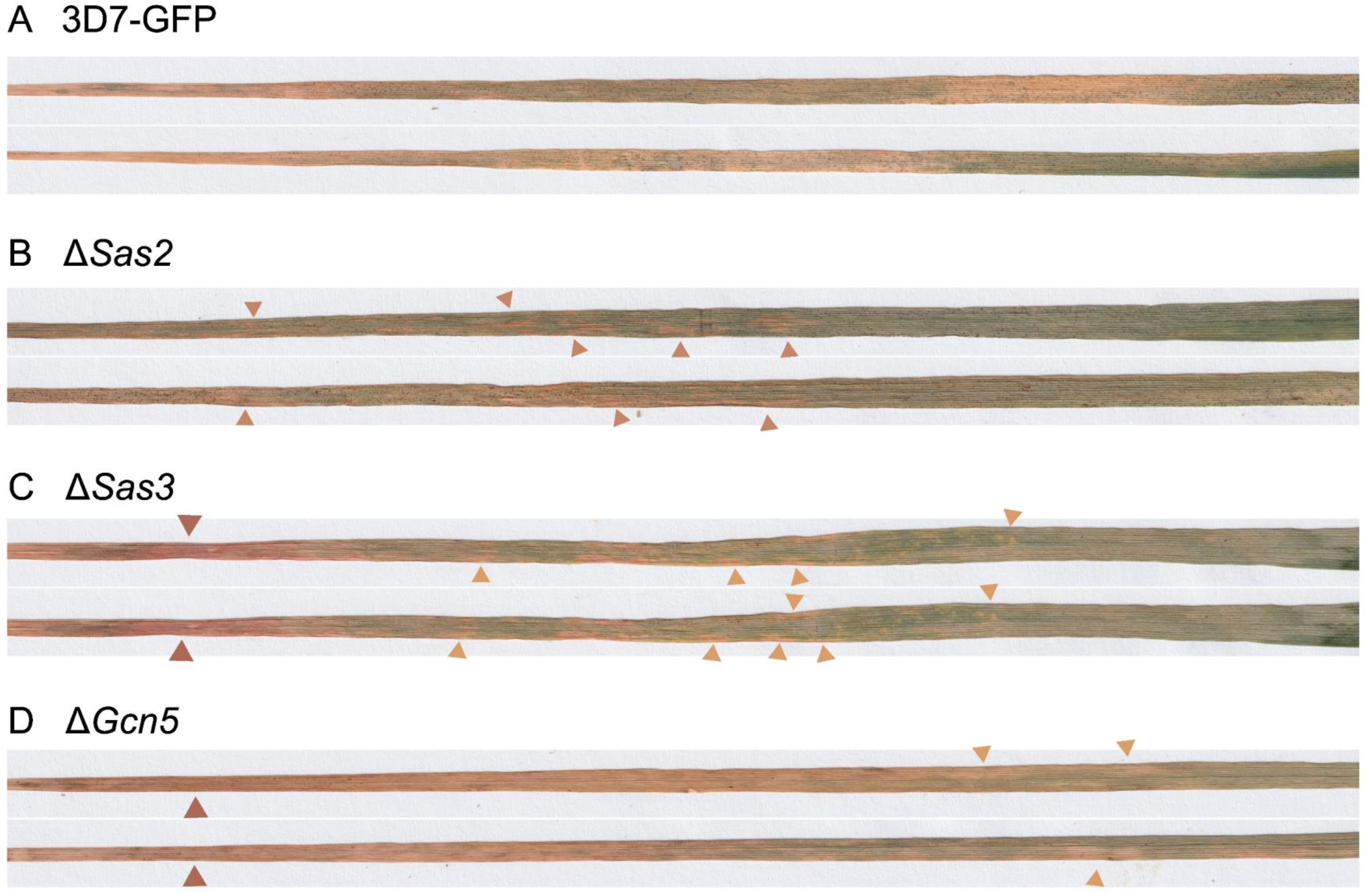
Δ*Sas2*, Δ*Sas3* and Δ*Gcn5* produce anormal macroscopic disease symptoms. Amplified pictures of leaves (length 14 cm) infected by 3D7-GFP (A), Δ*Sas2* (B), Δ*Sas3* (C) and Δ*Gcn5* (D) at 20 days post infection. Red/orange/yellow symptoms produced in the wheat cultivar Runal by the mutants are indicated with colored arrows. Note that no pycnidia were produced by Δ*Sas3* and Δ*Gcn5* in the spots with red/orange/yellow symptoms.

**Figure S7.**
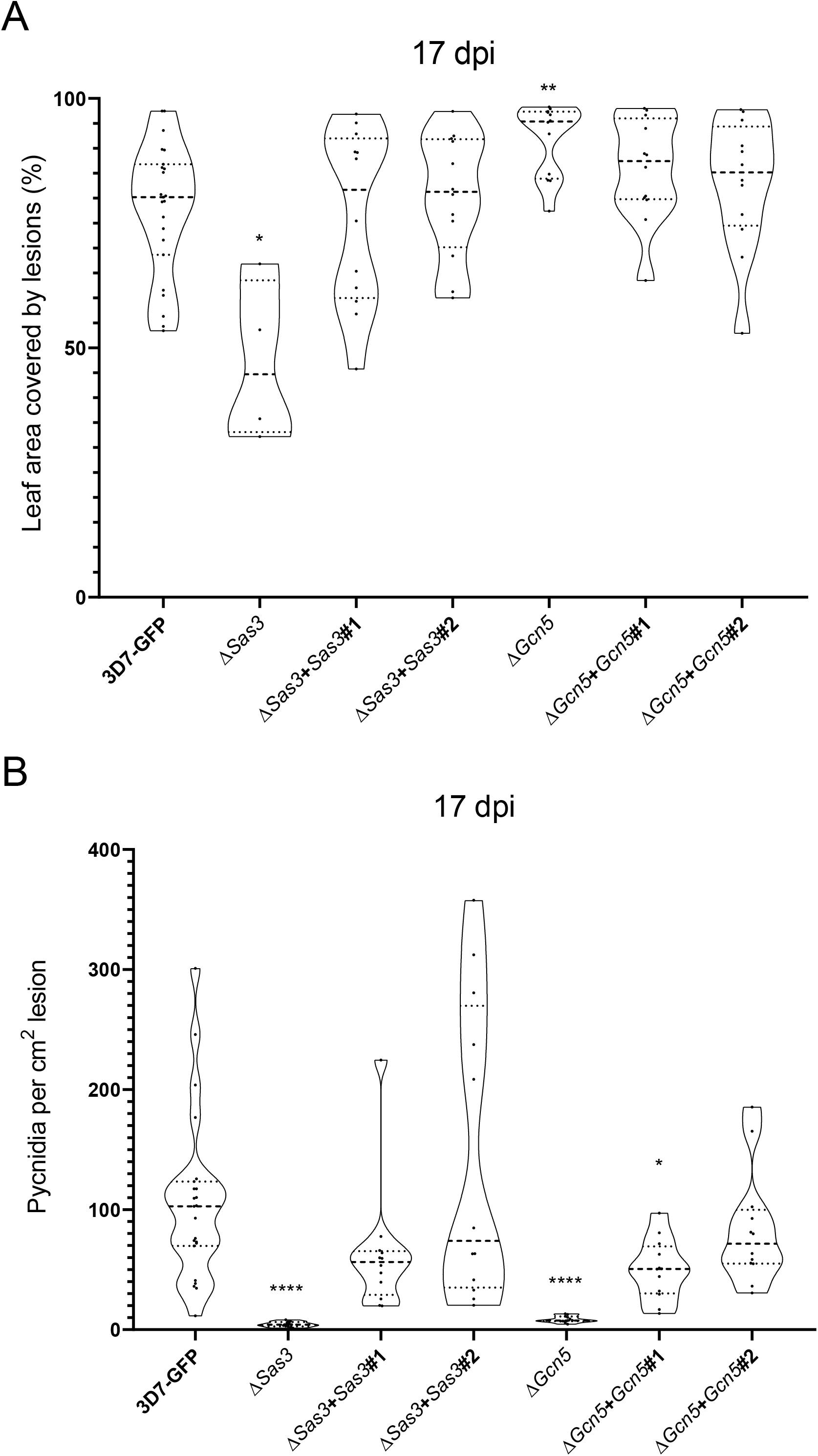
Complementation of Δ*Sas3* and Δ*Gcn5* recover the virulence phenotype. Percentage of leaf area covered by lesions (A) and pycnidia per cm^2^ of lesion (B) at 17 days post infection (dpi). Dashed lines represent the median, dotted lines represent first and third quartiles and black dots represent individual data points. Asterisks indicate statistically significant differences with 3D7-GFP according to the Kruskal-Wallis non-parametric statistical and posthoc uncorrected Dunn’s tests (* p < 0.05; ** p < 0.01; **** p < 0.0001).

**Figure S8.**
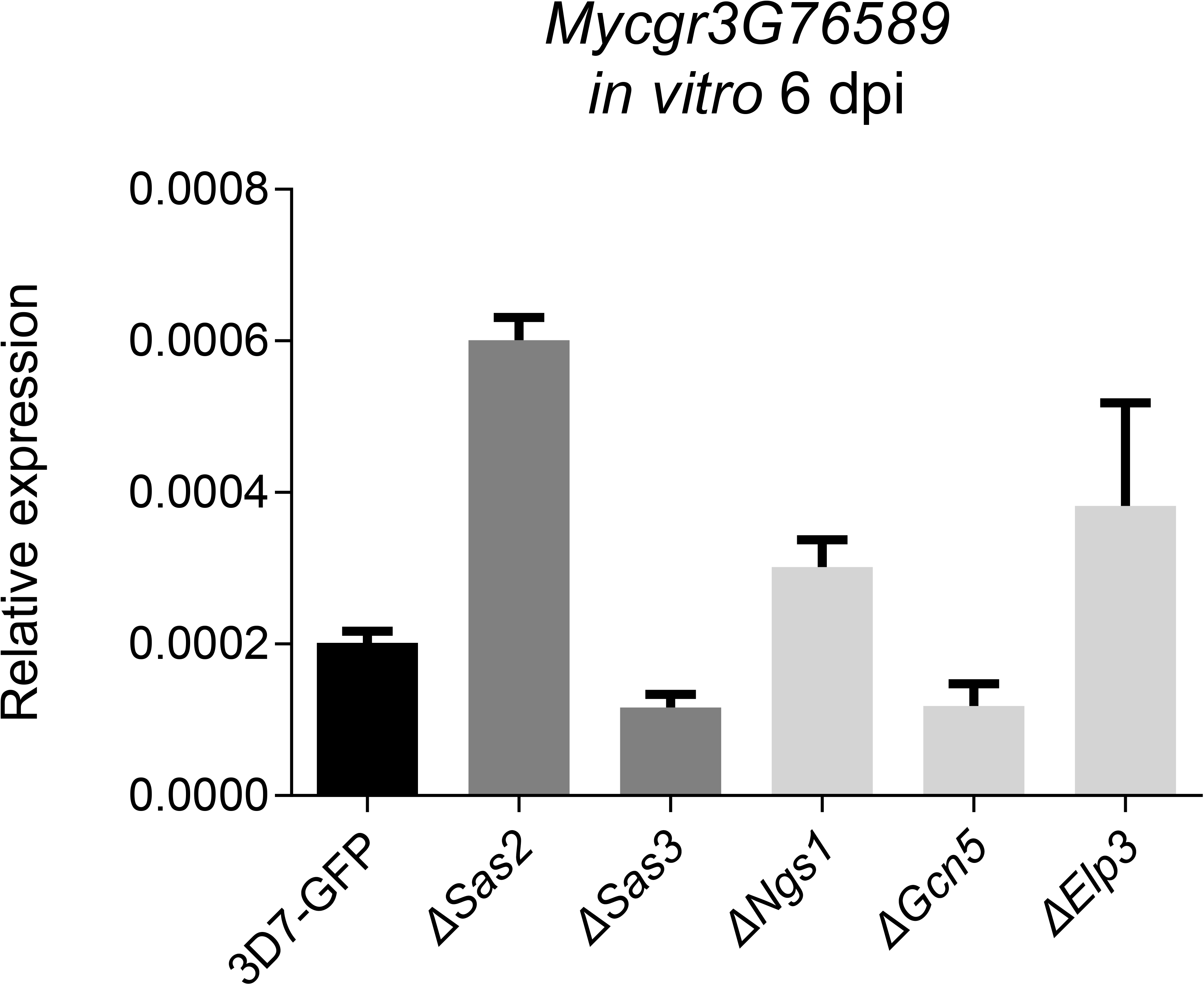
The expression pattern of *Mycgr3G76589* is not altered in lysine acetyltransferase (KAT) mutants under axenic conditions. Relative expression of *Mycgr3G76589* in 3D7-GFP and the KATs mutants grown on yeast-malt-sucrose agar (YMA) for 6 days. Tubulin and histone H3 were used as reference genes. Each bar corresponds to the average of 3 biological replicates. Error bars represent the standard error of the mean. No significant differences with 3D7-GFP according to the Kruskal-Wallis test were found (p < 0.05).

**Figure S9.**
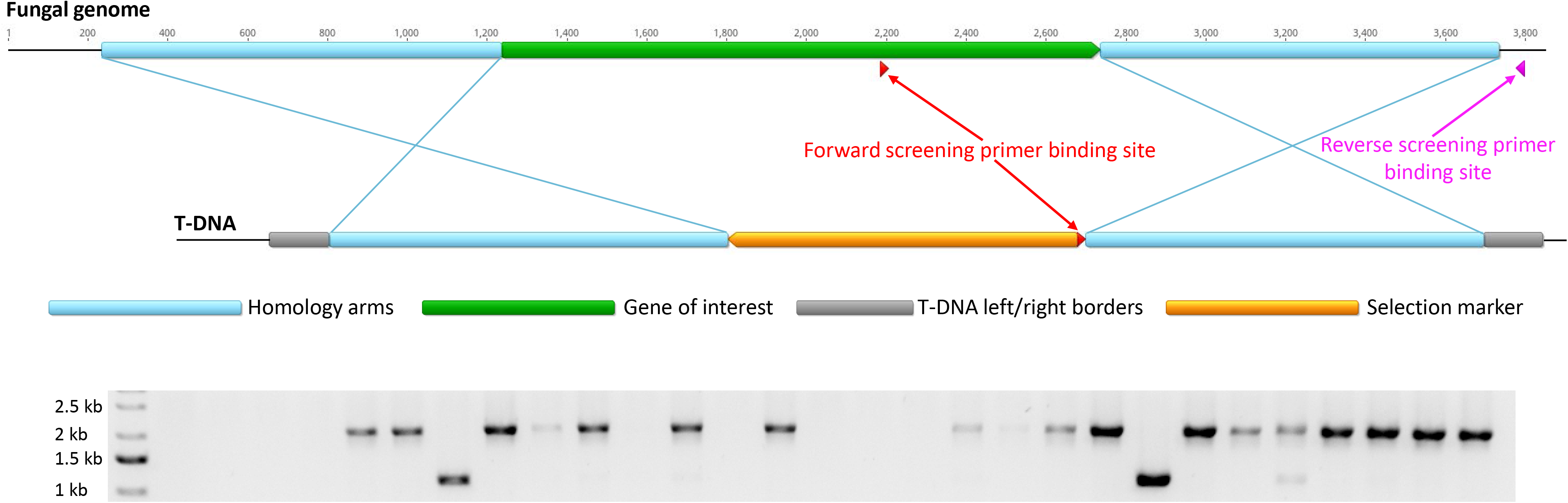
PCR screening strategy to identify gene deletion mutants. Top: Blue crosses represent homologous recombination events during fungal transformation. The forward screening primer binding sites within the gene of interest and the T-DNA are identical. Sizes of different parts do not correspond to a specific construct used in this study. Bottom: Example screening results for Δ*Elp3* mutant lines using liquid cultures directly added to the reaction. PCRs using forward and reverse screening primers yield distinct amplicons for native genes (1967 bp) and disrupted genes (1058 bp). Each lane represents a different mutant line.

## Notes

### Competing Interest Statement

The authors have declared no competing interest.

